# Ancient gene duplications, rather than polyploidization, facilitate diversification of petal pigmentation patterns in *Clarkia gracilis* (Onagraceae)

**DOI:** 10.1101/2021.04.26.441548

**Authors:** Rong-Chien Lin, Mark D. Rausher

## Abstract

It has been suggested that gene duplication and polyploidization create opportunities for the evolution of novel characters. However, the connections between the effects of polyploidization and morphological novelties have rarely been examined. In this study, we investigated whether petal pigmentation patterning in an allotetraploid *Clarkia gracilis* has evolved as a result of polyploidization. *C*. *gracilis* is thought to be derived through a recent polyploidization event with two diploid species, *C*. *amoena huntiana* and an extinct species that is closely related to *C*. *lassenensis*. We reconstructed phylogenetic relationships of the *R2R3*-*MYB*s (the regulators of petal pigmentation) from two subspecies of *C*. *gracilis* and the two purported progenitors, *C*. *a*. *huntiana* and *C*. *lassenensis*. The gene tree reveals that these *R2R3*-*MYB* genes have arisen through duplications that occurred before the divergence of the two progenitor species, i.e., before polyploidization. After polyploidization and subsequent gene loss, only one of the two orthologous copies inherited from the progenitors was retained in the polyploid, turning it to diploid inheritance. We examined evolutionary changes in these *R2R3*-*MYB*s and in their expression, which reveals that the changes affecting patterning (including expression domain contraction, loss-of-function mutation, *cis*-regulatory mutation) occurred after polyploidization within the *C*. *gracilis* lineages. Our results thus suggest that polyploidization itself is not necessary in producing novel petal color patterns. By contrast, duplications of *R2R3*-*MYB* genes in the common ancestor of the two progenitors have apparently facilitated diversification of petal pigmentation patterns.

## Introduction

Since the seminal work of Ohno (1970), duplicated genes have generally been thought to provide important material for the origin of evolutionary novelties. A duplicate gene copy can contribute to genetic and morphological diversification by evolving new gene functions (neofuctionalization), while the other copy can maintain the ancestral function (Zhang 2003; Rensing 2014). In addition, whole genome duplication (WGD) often leads to extensive changes in gene expression, which can potentially produce novel traits (Wang et al. 2012; Shi et al. 2015).

WGD events (involving either allo- or auto-polyploidization) have been common in the evolution of angiosperms (De Bodt et al. 2005; Flagel and Wendel 2009; Rensing 2014). Because angiosperms are the most species-rich group of plants and exhibit a great diversity of morphological and physiological traits, it seems likely that polyploidization has facilitated diversification and speciation in this group. However, the direct connections between the effects of polyploidization and morphological characters are in general unclear, largely because there are few studies that have attempted to ascertain how divergent copies of duplicated genes affect plant development. One impediment to such studies is that gene duplicates often become lost or silenced over short evolutionary timescales (Lynch and Conery 2000, 2003). It is thus difficult to establish whether diverged paralogs represent duplicate copies created by WGD or copies produced by tandem or segmental duplication after WGD and gene loss. This distinction is important because in the former situation, WGD actually provides the raw material for evolutionary novelty, whereas the latter situation is not a direct result of WGD.

This difficulty can be overcome by examining a recent polyploidization event in which the parental species are identifiable, and by following the inheritance, modification, loss or duplication of individual parental-species gene copies in the polyploid. Furthermore, if the effects of these gene copies on the phenotype can be determined, it should be possible to determine whether and how WGD contributes to the evolution of novel morphological traits. Here we adopt this approach to examine the effects of polyploidization on the evolution of novel petal pigment pattern elements in the genus *Clarkia* (Onagraceae).

In this study, we examine the allotetraploid *C*. *gracilis* (Piper) A. Nelson & J. F. Macbride and its two purported, diploid, progenitor species. *C*. *gracilis* is thought to be derived from *C*. *amoena huntiana* (Jeps.) H. Lewis & M. Lewis and an extinct species related to *C*. *lassenensis* (Eastw.) H. Lewis & M. Lewis and *C*. *arcuata* (Kellogg) A. Nelson & J. F. Macbride (Abdel-Hameed & Snow 1968, 1972), which have similar floral color patterns. We chose *C*. *lassenensis* to represent the extinct parental species, because a relatively better chromosome pairing was observed in *C*. *lassenensis* x *C*. *grailis* triploids than that in *C*. *arcuata* x *C*. *grailis* triploids (Hakansson 1946, cited by Abdel-Hameed & Snow 1972).

Although both of the two progenitor species have a pink petal background, they differ in floral color pattern. *C*. *a*. *huntiana* petals have red central spots (figs. 1A and 1F). By contrast, petals of *C*. *lassenensis* have red basal spots and narrow white bands above the spots (figs. 1B and 1G).

**Fig. 1.**
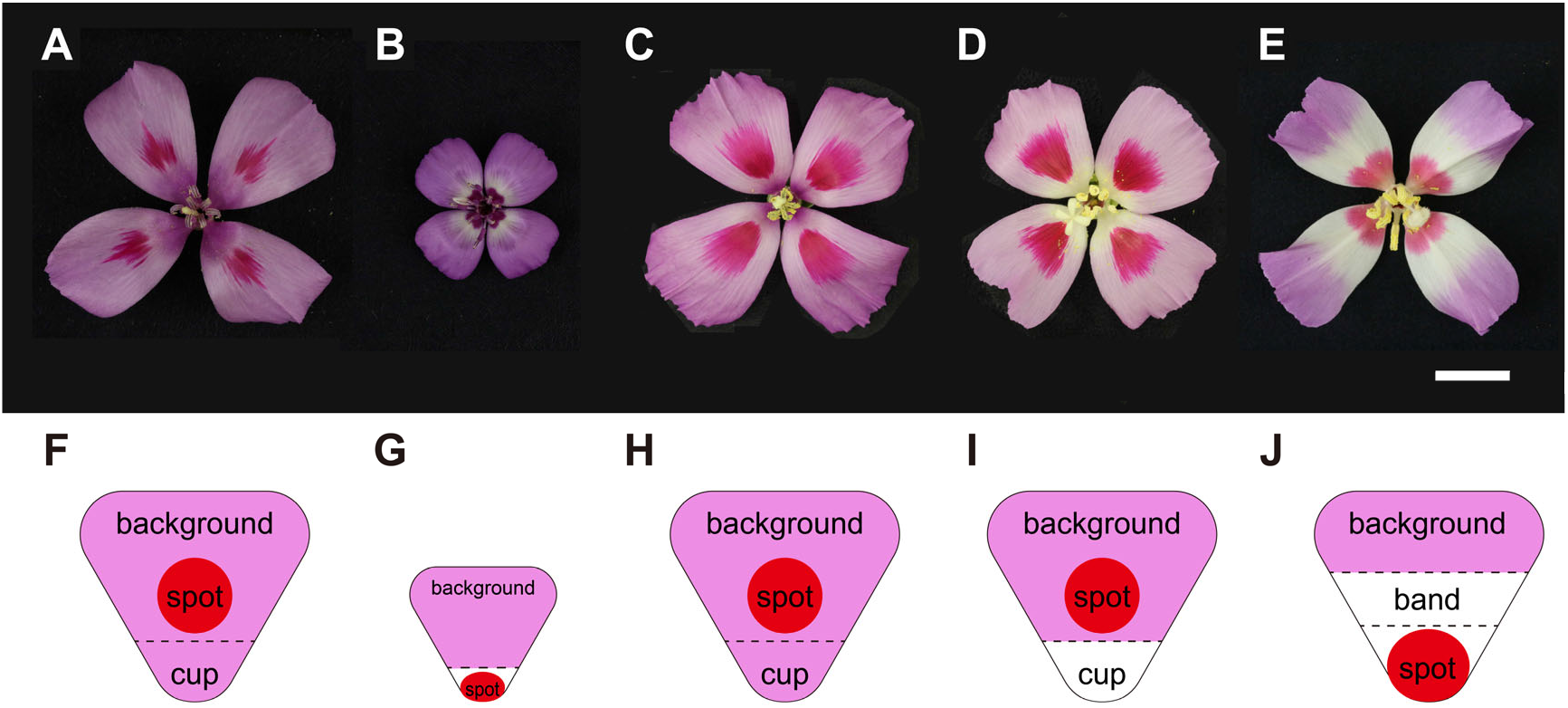
Flowers of the *Clarkia* (sub)species used in this study. (*A*) *C*. *amoena huntiana*; (*B*) *C*. *lassenensis*; (*C*) pink-cupped *C*. *gracilis sonomensis*; (*D*) white-cupped *C*. *g*. *sonomensis*; (*E*) *C*. *g*. *albicaulis*; (*F*-*J*) The elements of petal pigmentation patterns of these flowers. The scale bar indicates 15 mm.

The tetraploid *C*. *gracilis* has four named subspecies, two of which were included in this study. Both have color patterns that differ from those of the two parental species, as well as from each other, and thus represent the evolution of novelty. One is *C*. *g*. *sonomensis* that typically has petals with a pink background and red central spots (figs. 1C and 1H), while one of its variants has a basal petal region lacking pigmentation (i.e., “white cup”) (figs. 1D and 1I). The other subspecies, *C*. *g*. *albicaulis*, has a similar pink petal background, but has a basal spot and a large unpigmented (white) band in the middle of each petal (figs. 1E and 1J).

Previous studies have demonstrated that in *C*. *g. sonomensis*, each of the distinctive pattern elements (background, spot, cup) is controlled by different sets of *R2R3-MYB* transcriptional regulators (hereafter “*MYB*”) (Martins et al. 2017; Lin and Rausher 2021). *MYB1* regulates spot formation, while *MYB6*, *MYB11* and *MYB12* control background pigmentation (hereafter “background *MYB*s”), including presence/absence of the white cup (fig. 2). The protein products of these *MYB*s form complexes with bHLH and WDR proteins to activate the enzyme-coding genes in the anthocyanin biosynthetic pathway (Ramsay and Glover 2005; Xu et al. 2015). Because a single *bHLH* or *WDR* gene has broader expression domains and influences more characters than an individual *MYB* gene, the latter is largely responsible for tissue-specific/pattern-specific expression of anthocyanin pigments (Ramsay and Glover 2005; Albert et al. 2011; Streisfeld and Rausher 2011).

**Fig. 2.**
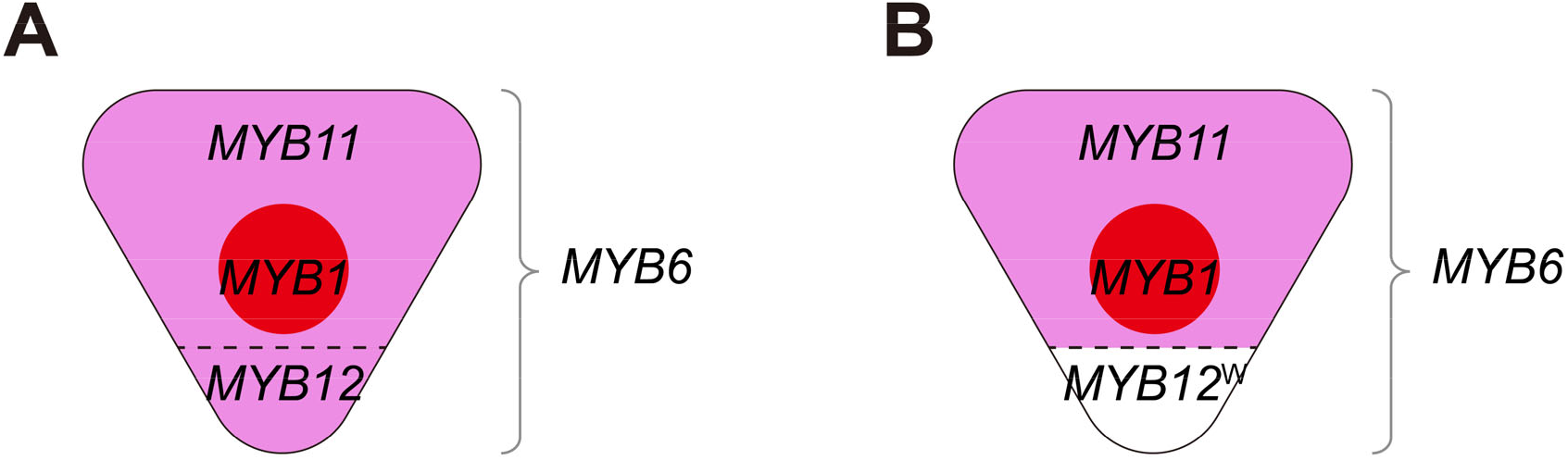
Schematic portrayal of expression domains of *R2R3*-*MYB* genes controlling color pattern elements in *Clarkia gracilis sonomensis*. *MYB1* controls spot formation. *MYB6* is expressed throughout the petal. *MYB11* is expressed in the distal petal region above the cup, whereas *MYB12* is expressed in the cup. (*A*) In conjunction with *MYB6*, *MYB11* and *MYB12* produce pigmentation in the distal and cup regions, respectively. (*B*) In individuals with a white cup, a nonfunctional *MYB12*^W^ is expressed in the cup (Lin and Rausher 2021).

Because of their central role in regulating color pattern elements, we undertook an examination on the evolution of these *MYB* genes in *C. gracilis* and its progenitors to determine whether polyploidization had direct effects on how these genes influenced pattern evolution. Specifically, we ask how changes in copy number, functionality or expression patterns of these genes contributed to the evolution of color pattern elements.

The effects of polyploidization on these *MYB* genes could directly affect pattern evolution by three distinct processes:

### Process 1

Polyploidization combines *MYB* genes controlling disparate pattern elements from the two parental species, creating a new pattern that is a combination of elements from the progenitors. For example, in *C. g. albicaulis*, the basal spot might be produced by the copy of *MYB1* controlling the basal spot in *C. lassenensis*, while the petal background pigmentation (including the white band) might be controlled by *MYB6*, *MYB11* and *MYB12* inherited from *C. a. huntiana.* Similarly, the white cup in *C. g. sonomensis* may reflect inheritance of *MYB* genes controlling the white cup in *C. lassenensis*, while the central spot may reflect inheritance of *MYB1* from *C. a. huntiana*.

### Process 2

Polyploidization results in two copies of orthologous *MYB* genes from the progenitors, which allows for subsequent neofunctionalization or subfunctionalization to produce new pattern elements. For example, the large white band in the middle of the *C*. *g*. *albicaulis* petal, which is lacking in both progenitors, may reflect neofunctionalization or subfunctionalization of duplicate copies of petal background *MYB*s inherited from the progenitors.

### Process 3

Through interactions between *MYB* genes from the two parental species, novel patterns that were not present in either parent can be generated. This type of interaction could also explain the central white band in *C*. *g*. *albicaulis*.

The alternative to a direct effect of WGD on the evolution of color patterns in *C. gracilis* is that pattern changes are caused by mutations affecting the *MYB* genes that could have produced the same change in one of the progenitor species. One example would be if the basal spot in *C*. *g*. *albicaulis* resulted from a mutation in the copy of *MYB1* inherited from *C*. *a*. *huntiana*, which produces a central spot in *C*. *a*. *huntiana*, rather than from inheritance of the copy of *MYB1* from *C. lassenensis*, which makes *C. lassenensis* basal-spotted. Another would be if the loss of pigmentation in the white cup of *C*. *g*. *sonomensis* represents an independent mutation in the background *MYB* gene responsible for pigmentation in the cup region rather than from inheritance of the background *MYB*(s) controlling the white cup in *C*. *lassenensis*. Yet another would be if the white band in *C*. *g*. *albicaulis* resulted from a mutation in a background *MYB* rather than from an interaction between genes inherited from the two progenitors. In each of these cases, polyploidization would not have been necessary in order for the changes to have evolved.

## Results

### *Identification of R2R3-MYB*s

Martins et al. (2017) demonstrated that in *C*. *gracilis*, *CgMYB1* is responsible for initiating spot formation early in the flower-bud development. Two different alleles (*CgMYB1C* and *CgMYB1B*) at this locus determine whether spotting is central (as in *C*. *g*. *sonomensis*) or basal (as in *C*. *g*. *albicaulis*). Lin and Rausher (2021) showed that three *R2R3-MYB* genes, *CgsMYB6*, *CgsMYB11* and *CgsMYB12*, are involved in petal background coloration in *C*. *g*. *sonomensis* (fig. 2). *CgsMYB6* is expressed throughout flower-bud development and everywhere in the petal, including the basal cup region. It activates all anthocyanin enzyme-coding genes except *CgsAns* (anthocyanidin synthase). *CgsMYB11* is expressed late in the development and activates *CgsAns*, which completes the expression of all enzyme-coding genes and allows pigments to form. This gene is not expressed, however, in the basal region of the petal (“cup”). Instead, pigmentation in the cup region is controlled by *CgsMYB12*. Like *CgsMYB11*, *CgsMYB12* is expressed late in development and activates *CgsAns* in this region. In the *C*. *g*. *sonomensis* individuals having the white cup, the copy of *CgsMYB12* is inactivated due to a premature stop codon.

From petal RNA of the progenitors *C*. *a*. *huntiana* and *C*. *lassenensis*, we cloned the full-length or nearly full-length copies of the four *R2R3-MYB* genes (supplementary fig. S1). Based on the primers used (supplementary table S1), these represent copies putatively orthologous to *CgMYB1*, *CgsMYB6*, *CgsMYB11*, and *CgsMYB12*. While cloning these genes from *C*. *g*. *albicaulis*, despite several attempts, we were unable to amplify the putative ortholog of *CgsMYB12* from this subspecies. Consistently, our transcriptome data also show that petal background transcriptome assemblies from *C*. *g*. *albicaulis* only reveal *MYB6* and *MYB11*, each of which has one copy (supplementary table S2; supplementary methods S1).

Because each of the two diploid progenitors expresses four *R2R3*-*MYB* genes, we would expect the two subspecies of *C*. *gracilis* to express eight different copies if there had been no gene loss or gene silencing following polyploidization. However, our recovery of at most four copies suggests that after polyploidization, four of these copies – one copy of each of the four paralogs in the progenitors – have either been lost, have been strongly downregulated, or have evolved sufficiently in sequence that they are no longer amplified by the primers used. Because petal background transcriptome assemblies from *C*. *g*. *albicaulis* and *C*. *g*. *sonomensis* (Lin and Rausher 2021) reveal only one copy of each of the four paralogs, the second copy of each gene has most likely been lost or downregulated in *C*. *gracilis*.

### *Phylogenetic relationships of R2R3-MYB*s

The reconstructed maximum-likelihood gene tree confirms the *MYB* copies in *C. gracilis* are orthologs of the genes identified in the progenitors (fig. 3). In particular, these genes form four highly-supported clades, each containing one copy from each of the four examined (sub)species, except the *MYB12* clade, which lacks the copy from *C. g. albicaulis*. Moreover, the *MYB1*, *MYB11*, and *MYB12* genes form a clade separate from the *MYB6* clade, which suggests that the former genes are derived from each other through two rounds of duplication prior to the divergence of the two diploid progenitors. Duplication of the ancestral copy first gave rise to *MYB1* and the common ancestor of *MYB11* and *MYB12*, and then the second duplication gave rise to *MYB11* and *MYB12*.

**Fig. 3.**
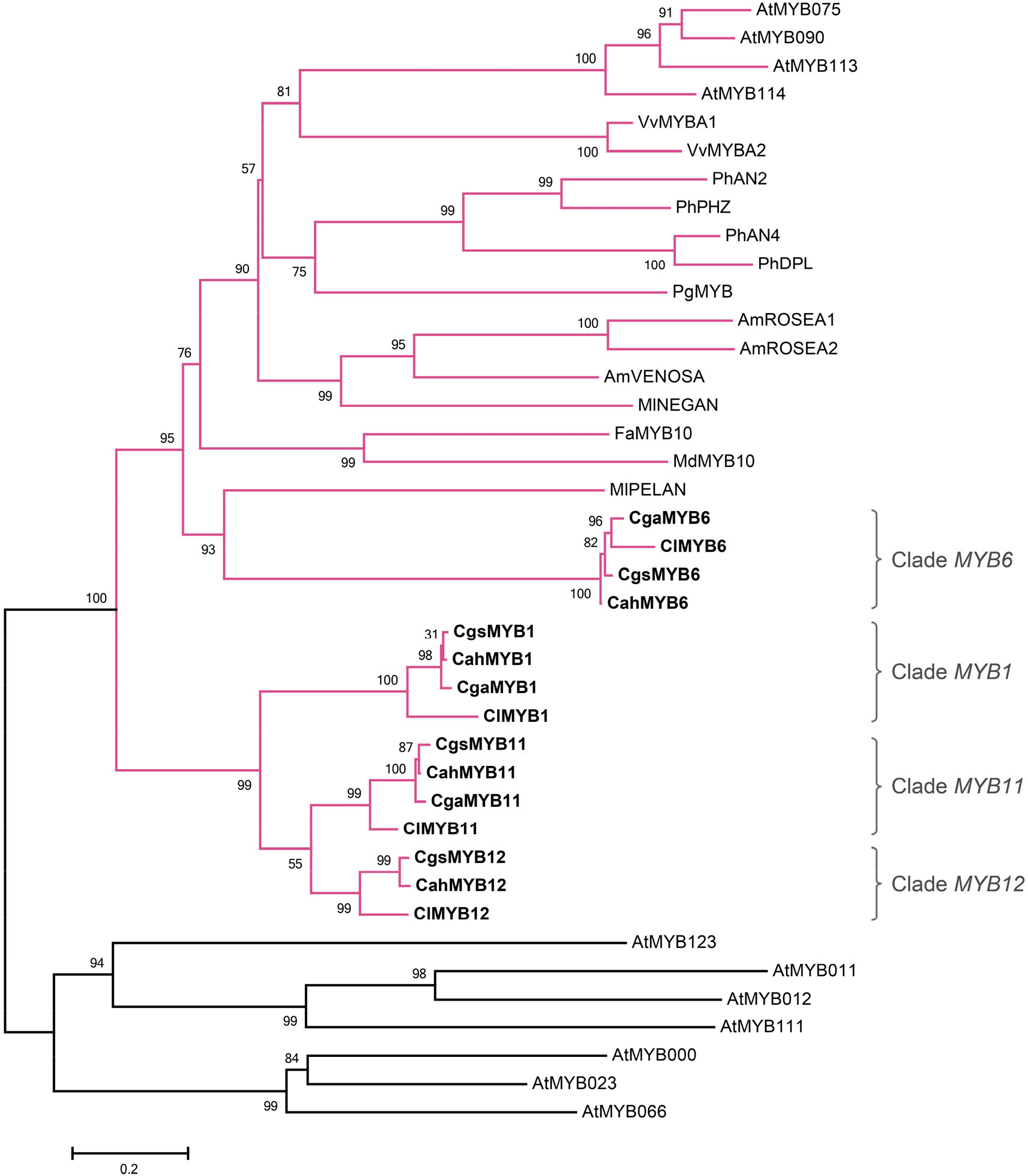
The maximum-likelihood gene tree of *R2R3-MYB*s. The clades containing the subgroup 6 *R2R3-MYB* genes, the regulators of the anthocyanin enzyme-coding genes, are shown in pink. The genes from the *Clarkia* (sub)species are shown in bold. Branch supports are shown as aRLT statistics. The *Arabidopsis thaliana* sequences were retrieved from TAIR (https://www.arabidopsis.org/): subgroup 5: *AtMYB123* (AT5G35550); subgroup 6: *AtMYB75* (AT1G56650), *AtMYB90* (AT1G66390), *AtMYB113* (AT1G66370) and *AtMYB114* (AT1G66380); subgroup 7: *AtMYB11* (AT3G62610), *AtMYB12* (AT2G47460) and *AtMYB111* (AT5G49330); subgroup 15: *AtMYB0* (AT3G27920), *AtMYB23* (AT5G40330) and *AtMYB66* (AT5G14750). Other sequences were retrieved from GenBank: *Antirrhinum majus AmROSEA1* (DQ275529), *AmROSEA2* (DQ275530), *AmVENOSA* (DQ275531); *Clarkia gracilis albicaulis CgaMYB1* (*CgMYB1B*, KX592431); *C*. *g*. *sonomensis CgsMYB1* (*CgMYB1C*, KX592432); *C*. *lassenensis ClMYB1* (KX592428); *Fragaria* x *ananassa FaMYB10* (EU155162); *Malus domestica MdMYB10* (EU518249); *Mimulus lewisii MlPELAN* (KJ011144), *MlNEGAN* (KJ011145); *Petunia* x *hybrida PhAN2* (AF146702), *PhAN4* (HQ428105), *PhDPL* (HQ116169), *PhPHZ* (HQ116170); *Punica granatum PgMYB* (KF841621); *Vitis vinifera VvMYBA1* (AB097923), *VvMYBA2* (AB097924).

Within the clades representing *MYB1*, *MYB11* and *MYB12*, the orthologs from *C*. *gracilis* are more closely related to the ortholog from *C*. *a*. *huntiana* than to the ortholog from *C*. *lassenensis*. This pattern, which has high statistical support (approximate likelihood ratio tests, aLRT), indicates that after polyploidization, it was the orthologs of each of these genes inherited from *C*. *lassenensis* that were lost or downregulated. By contrast, the copies of *MYB6* in *C*. *gracilis* are more similar to the copy from *C*. *lassenensis* than to the copy from *C*. *a*. *huntiana*, indicating that in the tetraploid, it was the copy of this gene from *C*. *a*. *huntiana* that was lost or downregulated.

### *Expression domains of R2R3-MYB*s

The expression patterns of *MYB6* and *MYB11* in *C*. *g*. *albicaulis*, *C*. *a*. *huntiana*, and *C*. *lassenensis* (fig. 4) are consistent with what has been previously reported for *C*. *g*. *sonomensis* (Lin and Rausher 2021). In these three (sub)species, *MYB6* is expressed early during flower-bud development and remains expressed in all petal sections throughout bud maturation. *MYB11* is expressed late in development, and is only expressed in the pigmented petal background. Notably, in *C*. *g*. *albicaulis*, the expression of *MYB11* does not extend into the region of the white band in the middle of the petal.

**Fig. 4.**
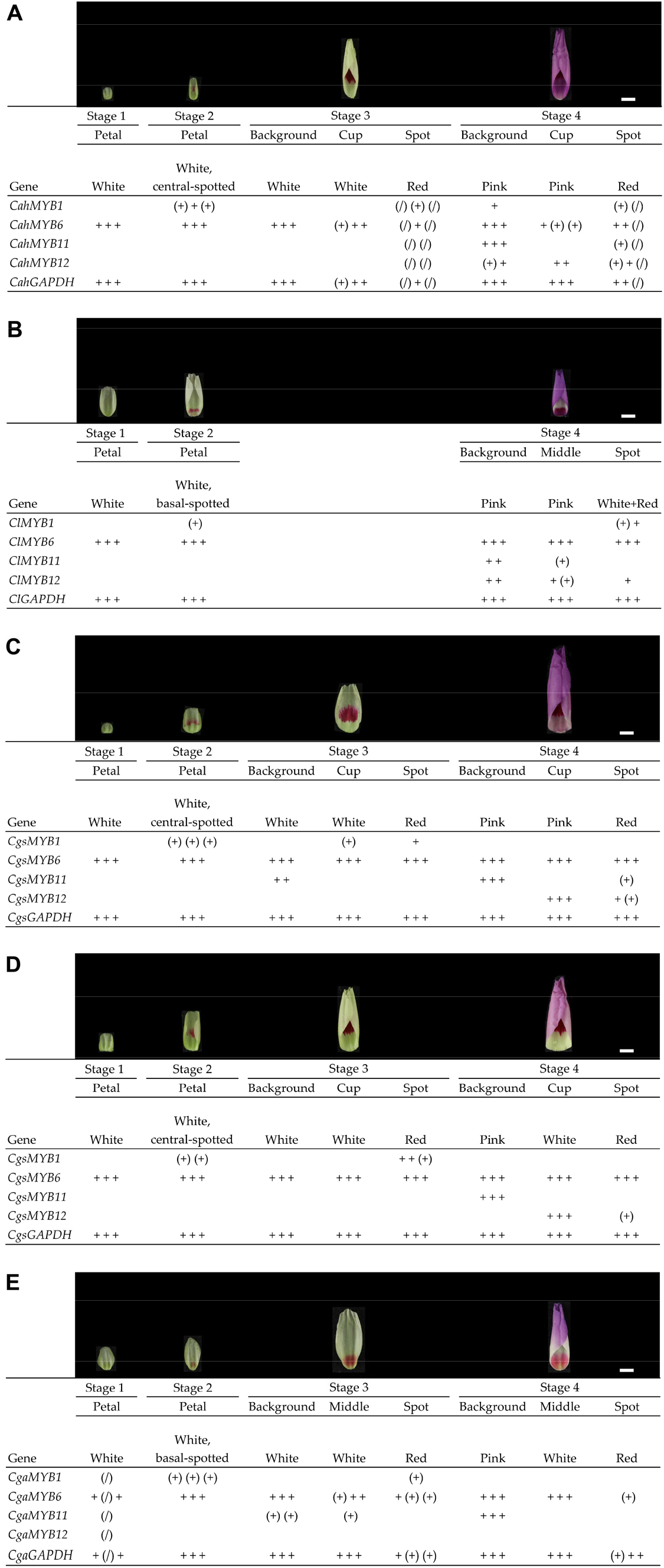
Expression patterns of *MYB1*, *MYB6*, *MYB11* and *MYB12* across the flower-bud development in (*A*) *Clarkia amoena huntiana*, (*B*) *C. lassenensis*, (*C*) pink-cupped *C*. *g*. *sonomensis*, (*D*) white-cupped *C*. *g*. *sonomensis* and (*E*) *C*. *g*. *albicaulis*. Based on the PCR-band brightness on the gels (see supplementary fig. S2 for gel photos), the expression levels were scored as “+”, “(+)”, and blank, respectively representing expressed, weakly expressed, and not expressed. A “+”, “(+)”, or blank represents a single plant. A “(/)” indicates a missing data point. A constitutively expressed gene *GAPDH* was included for cDNA quality control. Pictures above the columns designate the bud phenotypes. The scale bar indicates 5 mm.

By contrast, *MYB12* shows a more complex expression pattern (fig. 4). The timing of this gene’s expression is similar in *C*. *g*. *sonomensis* and the two diploid progenitors, being expressed late in the flower-bud development (Lin and Rausher 2021). However, its spatial expression domain differs from that of *MYB11*. In the two progenitors, *MYB12* is expressed throughout the petal, including basal (cup) region. Despite its expression in the cup region, this region is unpigmented in *C. lassenensis*, which may suggest that this gene is nonfunctional in *C. lassenensis*, although no evidence of frame shifts or premature stop codons was found in this gene, except a 27-bp insertion in Exon 3 (fig. S1D). By contrast, in *C*. *g*. *sonomensis*, *MYB12* is only expressed in the basal (cup) region, allowing pigmentation of that region (Lin and Rausher 2021). Expression of this gene was not detected in *C*. *g*. *albicaulis*, which, along with absence of *MYB11* expression in the middle portion of the petal, is consistent with absence of background pigmentation in the central and basal portions of the *C*. *g*. *albicaulis* petal.

Finally, *MYB1* expression in all four (sub) species (fig. 4) appears to be consistent with the pattern previously reported for *C*. *g*. *sonomensis* (Martins et al. 2017): its expression is limited to spots. We have found that in *C*. *a*. *huntiana*, *MYB1* expression was detected in a background sample at Stage 4 (fig. 4A). Given that this happened in only one out of three samples, we suspect that this result may reflect contamination of that sample. In *C*. *g*. *sonomensis*, however, we also found that it is expressed at appreciable levels in the cup region (fig. 4C). We suspect that this reflects expression in the small basal spot at the most proximal part of the cup region. This spot appears phenotypically similar to the central spot, being red rather than pink, suggesting they are activated by the same *MYB* copy.

In *C*. *g*. *albicaulis* and *C*. *lassenensis*, *MYB1* expression is restricted to the region of the basal spots. While this pattern suggests that spots in these two (sub)species are homologous, the gene tree indicates that this is not the case. Specifically, the basal spot in *C*. *g*. *albicaulis* is produced by a copy of *MYB1* that is more similar to, and thus inherited from, the central-spotted *C*. *a*. *huntiana* (fig. 3). This pattern implies that the basal position of the spot in *C*. *g*. *albicaulis* evolved independently after polyploidization. In particular, Martins et al. (2017) demonstrated that this shift in spot position was caused by a mutation in the *cis*-regulatory region of *CgMYB1*. Because this mutated copy in *C*. *g*. *albicaulis* was derived from the copy of *MYB1* inherited from the central-spotted *C*. *a*. *huntiana*, the basal position of the spot represents convergence on the basal-spotted phenotype exhibited by *C*. *lassenensis* rather than homology.

## Discussion

### A Model for the evolution of petal color patterns in Clarkia gracilis

The results reported here suggest a model for the evolution of petal color patterns in the tetraploid *C*. *gracilis* (fig. 5). This model makes the following assumptions: (1) pigmentation in the petal background requires expression of a functional *MYB6*, which activates all anthocyanin enzyme-coding genes except *Ans*, and either *MYB11* or *MYB12*, which activate *Ans* (Lin and Rausher 2021); (2) *MYB1* activates at least *Ans* in the regions where spots form.

**Fig. 5.**
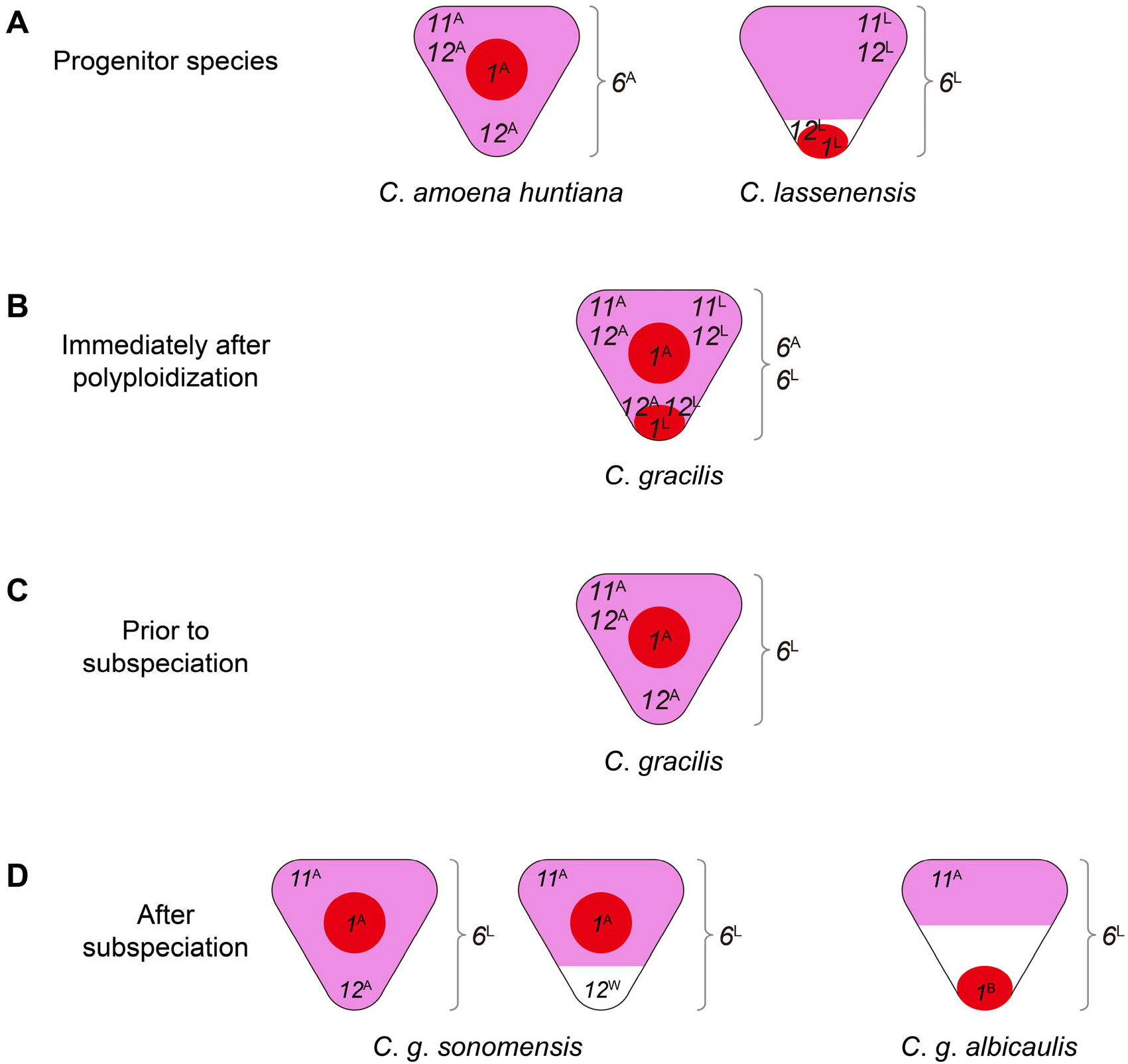
A model for the evolution of petal pigmentation patterning in *Clarkia gracilis*. (*A*) *R2R3-MYB* genes in the progenitor species of *C*. *gracilis*, *C*. *amoena huntiana* and *C*. *lassenensis*. *R2R3-MYB* genes are designated by number and letter: *1*, *MYB1*; *6*, *MYB6*; *11*, *MYB11*; *12*, *MYB12* and ^A^, the copy from *C*. *a*. *huntiana*; ^L^, the copy from *C*. *lassenensis*. The positions of the numbers indicate the expression domains of the *R2R3-MYB* genes. *6* (*MYB6*) is shown beside the petal because it is expressed throughout the whole petal. Colored areas indicate regions in which pigmentation is visible: red for spot formation, pink for background pigmentation, white for unpigmentation. (*B*) Immediately after polyploidization, genes from the two progenitors were all combined in the ancestor of *C*. *gracilis*. (*C*) Four changes occurred prior to subspeciation: gene loss/downregulation of *MYB1*^L^, *MYB6*^A^, *MYB11*^L^, and *MYB12*^L^. (*D*) Five changes occurred after subspeciation: expression domain contraction of *MYB12*^A^ in *C*. *g*. *sonomensis*; generating *MYB12*^W^ by a loss-of-function mutation in *C*. *g*. *sonomensis*; gene loss/downregulation of *MYB12*^A^ in *C*. *g*. *albicaulis*; expression domain contraction of *MYB11*^A^ in *C*. *g*. *albicaulis*; generating *MYB1*^B^ by a *cis*-regulatory mutation in *C*. *g*. *albicaulis*.

In the progenitors (fig. 5A), *MYB1* is expressed in regions that become spots, primarily centrally in *C*. *a*. *huntiana* (*MYB1*^A^) and basally in *C*. *lassenensis* (*MYB1*^L^). *MYB6*^A^ (in *C*. *a*. *huntiana*) and *MYB6*^L^ (in *C*. *lassenensis*) are expressed throughout the petal. The *MYB11* copies in *C*. *a*. *huntiana* and *C*. *lassenensis* (*MYB11*^A^ and*MYB11*^L^, respectively) are expressed in the distal regions of the petal, promoting, in conjunction with *MYB* 6, background pigmentation in those regions. Pigmentation in the basal (cup) region in *C*. *a*. *huntiana* is controlled by *MYB12*^A^ that is expressed in both distal and basal regions. Similarly, in *C*. *lassenensis*, *MYB12*^L^ is expressed throughout the petal, but there is a basal petal area that lacks any pigmentation. Because *MYB11*^L^ is not expressed in this region, a functional copy of *MYB12*^L^ would be required for pigmentation. Thus, one explanation for the white cup in *C*. *lassenensis* is that *MYB12*^L^ is nonfunctional. However, we cannot rule out the possibility that some other factor (such as an inhibitor) is preventing pigmentation in this area.

Immediately after polyploidization (fig. 5B), *C*. *gracilis* presumably expressed all eight copies of these genes. The entire petal background would have been pigmented because there would have been functional *MYB6*^A^ and *MYB6*^L^, and either *MYB11*s or *MYB12*s expressed in all petal regions. In particular, the cup region would be pigmented because *MYB12*^A^ was functional. Expression of *MYB1*^A^ would presumably have produced a central spot and *MYB1*^L^ a basal spot. The petal would thus have had a pink background with two spots (fig. 5B).

The phylogenetic evidence indicates that prior to subspeciation (fig. 5C), *MYB1*^L^, *MYB6*^A^, *MYB11*^L^ and *MYB12*^L^ were either lost or downregulated. We infer that *MYB1*^L^, the *MYB1* copy from *C*. *lassenensis*, was lost or downregulated because the *MYB1* copies in the two *C*. *gracilis* subspecies are more similar to *MYB1*^A^, the *C*. *a*. *huntiana* copy. This inference is also consistent with the central location of the spot in *C*. *g*. *sonomensis*. Since functional copies of all four genes remained and were presumably expressed, the petals of the common ancestor of the two *C*. *gracilis* subspecies had a pink background throughout the petal and a single central spot (fig. 5C).

After subspeciation (fig. 5D), *C*. *g*. *sonomensis* underwent two changes: (1) the expression domain of *MYB12*^A^ contracted, such that it is expressed only in the proximal (cup) region; (2) a new, loss-of-function mutation occurred in that gene, producing *MYB12*^W^ (Lin and Rausher 2021), creating the pink/white cup polymorphism. In *C*. *g*. *albicaulis*, *MYB12*^A^ became downregulated or was nonfunctionalized or deleted. Additionally, the expression domain of *MYB11*^A^ in *C*. *g*. *albicaulis* contracted to just the most distal region of the petal. This contraction produced a white band in the middle of the petal, where neither *MYB11*^A^ nor *MYB12*^A^ is expressed. We do not know, however, whether this contraction is due to a change in *MYB11*^A^ itself, or in upstream regulators or inhibitors.

Finally, based on the orthology of *MYB1* in the two *C*. *gracilis* subspecies, a *cis-* regulatory mutation in *MYB1*^A^ produced a new allele, *MYB1*^B^, which shifted its expression domain to the basal region (Martins et al. 2017). This allele became fixed in *C*. *g*. *albicaulis*, shifting the spot position from central to basal.

This model highlights the role of ancestral gene duplication prior to tetraploidization in facilitating the evolution of novel characters, specifically the white band in *C. g. albicaulis* and the white cup in *C. g. sonomensis.* Both of these novel pattern elements evolved because an ancestral duplication produced the paralogs *MYB11* and *MYB12.* After this duplication, their expression domains diverged, such that in the progenitor species, *MYB12* was expressed throughout the petal, whereas *MYB11* was not expressed in the cup region. This expression domain divergence was further enhanced in *C. g. sonomensis*, with *MYB12* expressed only in the cup region. This spatial differentiation allowed a functionally inactivating mutation in *MYB12* (*MYB12*^W^) to produce an unpigmented area in only part of the petal (the white cup) in *C. g. sonomensis.* Additionally, contraction of the *MYB11* expression domain in *C. g. albicaulis* created a region in the center of the petal in which neither *MYB11* nor *MYB12* was expressed, producing the white band.

### Effects of polyploidization on petal color patterns

In the model described above, there are nine evolutionary changes to *R2R3*-*MYB* genes in the polyploidy *C. gracilis*. Four of them occurred prior to subspeciation: (i) gene loss or gene downregulation of *MYB1*^L^; (ii) gene loss/downregulation of *MYB6*^A^ (iii) gene loss/downregulation of *MYB11*^L^; (iv) gene loss/downregulation of *MYB12*^L^. The others occurred after subspeciation: (v) *MYB12*^A^ expression domain contraction in *C*. *g*. *sonomensis*; (vi) a loss-of-function mutation in *MYB12*^W^ in *C*. *g*. *sonomensis*; (vii) gene loss/downregulation of *MYB12*^A^ in *C*. *g*. *albicaulis*; (viii) *MYB11*^A^ expression domain contraction in *C*. *g*. *albicaulis*; (ix) a *cis*-regulatory mutation in *MYB1*^B^ in *C*. *g*. *albicaulis*. Some of these changes (e.g., vi, viii, and ix) affect petal pigmentation patterning. However, none of the changes affecting patterning appear to be the direct result of polyploidization. Rather, they appear to be evolutionary changes within lineages of *C*. *gracilis* that occurred after one copy of each of the four *MYB*s had been lost or silenced – changes that could have occurred in a diploid species.

In particular, for the two novel phenotypic changes that occurred in *C. gracilis*, we found no evidence for any of the three processes involving direct effects of polyploidization described in Introduction. One phenotypic change is a candidate for Process 1, the combining of *MYB* genes for different elements of the two progenitors to create a new pattern: the combination of a central spot (present in *C. a. huntiana*) with a white cup (present in *C. lassenensis*) in *C. g. sonomensis*. However, we have shown that the white cup in this species is derived from a mutation in the copy of *MYB12* inherited from *C. a. huntiana* as well.

The second phenotypic change – the presence of a white band in the center of the *C. g. albicaulis* petals – is a candidate for either Process 2 or 3. Process 2, which involves differentiation of the expression domain of two *MYB* orthologs, could not have occurred because only one copy of each paralog remained prior to subspeciation, which in turn occurred before evolution of the white band. Process 3, which involves interactions between *MYB* genes from different parents, could not have occurred because if it had, it presumably would have occurred at the time of polyploidization or soon thereafter, and we would thus expect both *C. gracilis* subspecies to exhibit the central white band. Since it does not appear in *C. g. sonomensis*, any gene interaction was likely not occurring at the time of subspeciation. Instead, the white band is found only in *C. g. albicaulis* and clearly results in a loss of expression (or gene loss) of *MYB12*^A^ and contraction of the domain of *MYB11*^A^ in that lineage.

One possible limitation of this study is that we have not characterized upstream regulators of the *MYB* genes, partly because the regulation of MYB-bHLH-WDR genes themselves is less understood (Xu et al. 2015). It is possible that an upstream regulator may have diverged in a way that would be a direct effect of WGD. For example, consider a regulator of *MYB1*. Although only one copy of this gene (*MYB1*^A^) is present in *C. gracilis*, it is possible that two copies of its regulator may be present, one inherited from each progenitor. Initially, the expression domain of both copies would produce a central spot. However, if the expression domain of one of the regulator copies shifted to the base of the petal, that would also cause a shift in the spot position, and could account for the basal spot in *C. g. albicaulis*. This would reflect Process 2 in Introduction. We know, however, that the basal spot position in *C. g. albicaulis* is cause by a *cis*-regulatory mutation in *MYB1*^A^ itself, ruling out this possibility.

Another possibility involves the expression domain contractions of *MYB11* and *MYB12* that occurred in *C. gracilis.* These contractions can be explained by changes in their regulators consistent with Process 2. For example, if *C. gracilis* retained both copies of a regulator of *MYB11*^A^, initially both would cause background pigmentation in the entire petal except for the cup region. If there was subsequent expression-domain subfunctionalization, however, the potential for formation of a white band would be present. Specifically, if the expression domain of one copy of the regulator contracted to just the distal portion of the petal, while the expression domain of the second copy contracted to the central portion of the petal, and this was followed by downregulation or loss of function of the second regulator copy, a white band would be produced. This scenario would be an example of Process 2 and would thus be a direct effect of polyploidization. However, other processes that are not direct effects can also produce a white band. For example, if one of the regulator is lost, the remaining regulator may undergo a contraction in expression domain to the distal portion of the petal. In this situation, there would be no *MYB* expressed in the central portion to activate *Ans*, complete the pathway, and cause pigments to be expressed. While at present we cannot distinguish between these two types of processes, the appearance of the novel white band in *C. g. albicaulis* is certainly consistent with processes that could operate regardless of whether WGD had occurred.

### Conclusions

While it has been suggested that polyploidization creates opportunities for the evolution of novel characters, this suggestion is based largely on the observation that morphological novelties (for example, floral forms; Zahn et al. 2005) have often arisen after polyploidization. However, there have been very few prior investigations that have attempted to determine whether the genome-combining effects of polyploidization itself are responsible for those changes. In this study, we provide evidence indicating that the evolution of novel phenotypes after tetraploidization in *C*. *gracilis* was likely not caused by the effects of polyploidization itself, but represent evolutionary changes that could have occurred if polyploidization had not taken place – the equivalent changes could have occurred in a diploid species. While it is dangerous to make generalizations based on one study, our results suggest that polyploidization itself may not contribute to trait diversification as much as is currently believed. Rather, ancient duplications of *R2R3*-*MYB* genes before polyploidization have apparently facilitated diversification of petal pigmentation patterns in *C*. *gracilis*. Our study also supports the common observation that evolutionary changes in floral pigmentation are accomplished primarily through modification of *R2R3-MYB* genes or their expression.

## Materials and Methods

### Plant growth

Methods for germination of the seeds of *C*. *a*. *huntiana*, *C*. *lassenensis*, and *C*. *g*. *albicaulis* (see supplementary table S3 for voucher information) and growth of these plants were described in Lin and Rausher (2021).

### Cloning of the R2R3-MYB genes

We amplified the coding regions of *MYB6*, *MYB11* and *MYB12* from *C*. *a*. *huntiana*, *C*. *lassenensis* and *C*. *g*. *albicaulis* with the primers listed in supplementary table S1. We also amplified *MYB1* from *C*. *a*. *huntiana* because the available sequence in GenBank is only 271-bp long (GenBank accession no. KX592430).

Total RNA of the collected/dissected petals (see supplementary fig. S2) was extracted using Spectrum Plant Total RNA Kit (Sigma-Aldrich, St. Louis, MO, USA). cDNA was synthesized following the methods described in Supporting Information Methods S4 in Lin and Rausher (2021). Amplification and sequencing of these four *R2R3*-*MYB* genes were conducted following Supporting Information Methods S5 in Lin and Rausher (2021). The sequences generated in this study were deposited at NCBI under GenBank accession numbers MT796894-MT796902.

### Phylogenetic analysis

The nucleotide sequences of *MYB1*, *MYB6*, *MYB11*, and *MYB12* from *C*. *a*. *huntiana*, *C*. *lassenensis*, *C*. *g*. *sonomensis*, and *C*. *g*. *albicaulis* and the subgroup 6 *R2R3-MYB*s from *Arabidopsis thaliana*, *Antirrhinum majus*, *Fragaria* x *ananassa*, *Malus domestica*, *Mimulus lewisii*, *Petunia x hybrida*, *Punica granatum*, and *Vitis vinifera* were aligned using MUSCLE (Edgar 2004). A maximum-likelihood phylogenetic tree was constructed using PhyML version 20120412 (http://www.atgc-montpellier.fr/phyml) (Guindon et al. 2010). The GTR+I+G (I = 0.080, G = 1.806) substitution model was used as determined based on the Akaike Information Criterion (AIC) by Smart Model Selection (SMS) version 1.8.1 (Lefort et al. 2017), which was integrated into PhyML. Clade support was estimated by approximate likelihood ratio tests (aLRT) based on a Shimodaira–Hasegawa-like procedure (Anisimova & Gascuel 2006).

### Semi-quantitative assessment of gene expression across flower-bud developmental stages

To examine the expression patterns of *MYB1*, *MYB6*, *MYB11*, and *MYB12*, we collected flower-buds from three plants each of *C*. *a*. *huntiana*, *C*. *lassenensis*, and *C*. *g*. *albicaulis*. The flower-buds were collected at four different stages that color appears in different pattern elements: (1) white petal; (2) central or basal spot appearing, depending on species; (3) central or basal spot well defined; (4) background and cup colors appearing. The larger petals (Stages 3 and 4) were dissected into sections as illustrated in supplementary fig. S2. For *C*. *lassenensis*, Stages 2 and 3 were combined into Stage 2 due to a small petal size. Samples of pink-cupped *C*. *g*. *sonomensis* (fig. 1C) and white-cupped *C*. *g*. *sonomensis* (fig. 1D) from Lin and Rausher (2021) (Types I and III, respectively) were also included for the comparison purpose.

Each of the cDNA samples prepared as described above was diluted to 2.5 ng/μl for semi-quantification of gene expression. PCR reactions were conducted using *Taq* DNA Polymerase (New England BioLabs, Ipswich, MA, USA) with the primers listed in supplementary table S1. The thermoprofile included: denaturation at 95 °C for 2 min, followed by 35 cycles of 95 °C for 30 s, 60 °C for 30 s, and 72 °C for 30 s, and final extension at 72 °C for 2 min. PCR products were visualized on 2% agarose gels. Gel photographs are shown in supplementary fig. S2. The brightness of PCR bands reflects the expression levels of the tested genes, which was scored as expressed (“+”), weakly expressed (“(+)”) or not expressed (blank). We also labeled the missing data as “(/)”.

## Supplementary Material

Supplementary tables S1, S2 and S3; Supplementary figures S1 and S2; Supplementary methods S1

## Acknowledgements

This work was supported by National Science Foundation Grant DEB1542387 to M.D.R. and by the Duke Biology grant to R.-C.L.. We are grateful to Talline Martins and Rancho Santa Ana Botanical Garden for providing seeds. We thank Duke University Greenhouse for plant care. We also thank members of the Rausher lab for constructive feedback.

## Notes

http://doi.org/10.5281/zenodo.4699974

### Competing Interest Statement

The authors have declared no competing interest.

