## Supplementary Material for "Ancient gene duplications, rather than polyploidization, facilitate diversification of petal pigmentation patterns in *Clarkia gracilis* (Onagraceae)"

**Table S1.** Primers used in this study.

**Table S2.** Anthocyanin genes identified from the transcriptomes of *Clarkia gracilis albicaulis*.

**Table S3.** Voucher information for the *Clarkia* plants used in this study.

**Fig. S1.** Alignment of *Clarkia R2R3-MYB* sequences (*A*) *MYB1*, (*B*) *MYB6*, (*C*) *MYB11* and (*D*) *MYB12* from *C. gracilis sonomensis*, *C. g. albicaulis*, *C. amoena huntiana* and *C. lassenensis*.

**Fig. S2.** Spatiotemporal expression of *MYB1*, *MYB6*, *MYB11* and *MYB12* in the petals of (*A*) *Clarkia amoena huntiana*, (*B*) *C. lassenensis*, (*C*) pink-cupped *C. gracilis sonomensis*, (*D*) white-cupped *C. g. sonomensis*, (*E*) *C. g. albicaulis*.

**Methods S1.** Transcriptomics

Table S1. Primers used in this study. [†]Primers from Martins *et al*. 2017. [‡]Primers from Lin and Rausher 2021. [a]Semi-qPCR primers used for *Clarkia lassenensis*.

**Primers for coding region sequencing**

Target species: *Clarkia amoena huntiana*

| Primer Name | Primer Sequence (5'-3') |
|---|---|
| cMYB1-F[†] | ATGAATAAGGTAGGACTTAGAAAGG |
| cMYB1-R[†] | TTAAAATATGTCATCAAAATAAAGCTCATCC |
| cMYB6-3F[‡] | TGCTACAGAAAGTCTAACGT |
| cMYB6-1R[‡] | ACCGCTGATTTATTTGAAACCCT |
| cMYB11-5F[‡] | AAAAACCAGAAGAAAACCCA |
| cMYB11-6R | AATTTCACAGTTTAATCATTTG |
| cMYB12-9F | AATGAAGGAAGGTCTAAGGAA |
| cMYB12-5R | CTACAACTGAAATATGTCGTGATC |

Target species: *Clarkia lassenensis*

| Primer Name | Primer Sequence (5'-3') |
|---|---|
| cMYB6-4F | CATGGGTGGTGTTCCTTGGA |
| cMYB6-5R | TTACAGAGAGTTATTCCACAGATCG |
| cMYB11-7F | AATGAAGGGAGAATTAAGGAAG |
| cMYB11-5R | TTACTTGGAAATATGATTCTACACA |
| cMYB12-11F | ATGCACGCTCTACAATAAAACG |
| cMYB12-6R | TTTAGTATAACGGCATAATTTCT |

Target species: *Clarkia gracilis albicaulis*

| Primer Name | Primer Sequence (5'-3') |
|---|---|
| cMYB6-3F[‡] | TGCTACAGAAAGTCTAACGT |
| cMYB6-1R[‡] | ACCGCTGATTTATTTGAAACCCT |
| cMYB11-3F | GTGAGTCGCATGGCTGT |
| cMYB11-2R[‡] | CACAGTTTAATCATTTGATTC |

**Primers for semi-quantitative assessment**

| Primer Name | Primer Sequence (5'-3') |
|---|---|
| cMYB1-gF[†] | TGGTCACTCATAGCAGGAAGA |
| cMYB1-Q1R | TCATCAACCAGTCCGACCAA |
| ClMYB1-1R[a] | GGGTTATACCCACTCTCAGTTCC |
| cMYB6-Q1F[‡] | GACGAACCCACCGAATCAAG |
| cMYB6-Q1R[‡] | CAGATCGTCCCAGTCCCATT |

| | |
|---|---|
| ClMYB6-Q1F[a] | CGGTGTCGGTCTTGGAGATA |
| ClMYB6-Q1R[a] | TCATCCCAATCCCATTTTCTGC |
| cMYB11-Q1F[‡] | GACAAGGAGGTGATGAATTGGT |
| cMYB11-Q1R[‡] | ATTCTACACATTCATGGAGTCGA |
| cMYB12-Q1F[‡] | GAGCAAGATTCGGTCAAAGT |
| cMYB12-1R[‡] | TATTCCGTTACAATGAGGCT |
| GAPDH-Q1F[‡] | GAGGCATCAGAGACCCACAT |
| GAPDH-Q1R[‡] | CACGACACGAGCTTCACAAA |

Table S2. Anthocyanin genes identified from the transcriptomes of *Clarkia gracilis albicaulis*. The transcriptomes were obtained from pink background and white band of the *C. g. albicaulis* petals, following the protocol described in supplementary methods S1. Gene expression levels were estimated as FPKM values by mapping reads to the transcriptome references of pink background and white band, separately.

| Contig ID | Gene | Expression (FPKM) | |
|---|---|---|---|
| | | Pink background | White band |
| *Pink background reference* | | | |
| RCL6_27068_c0_g1 | *Chs* | 1350.66 | 1138.03 |
| RCL6_21692_c0_g1 | *Chi* | 74.56 | 26.34 |
| RCL6_23311_c0_g4 | *F3h* | 45.94 | 50.03 |
| RCL6_27159_c0_g1 | *F3'h* | 15.98 | 27.28 |
| RCL6_23784_c0_g1 | *F3'5'h* | 649.53 | 381.47 |
| RCL6_23661_c0_g1 | *Dfr* | 757.58 | 121.86 |
| RCL6_26489_c1_g1 | *Dfr3* | 66.31 | 1140.31 |
| RCL6_26565_c4_g2 | *Ans* | 628.81 | 45.37 |
| RCL6_22269_c2_g4 | *Uf3gt* | 216.18 | 154.25 |
| RCL6_24035_c1_g1 | *MYB6* | 858.87 | 595.14 |
| RCL6_21823_c2_g1 | *MYB11* | 121 | 46.94 |
| RCL6_28257_c0_g1 | *bHLH1* | 39.6 | 34.93 |
| RCL6_20240_c0_g1 | *bHLH2* | 4.88 | 5.04 |
| RCL6_23159_c0_g2 | *WDR1* | 20.9 | 14.74 |
| RCL6_21868_c0_g1 | *WDR2* | 14.88 | 13.33 |
| *White band reference* | | | |
| RCL7_26363_c0_g2 | *Chs* | 1372.81 | 846.27 |
| RCL7_20945_c0_g1 | *Chi* | 108.08 | 25.05 |
| RCL7_20216_c0_g4 | *F3h* | 78.11 | 70.13 |
| RCL7_26446_c0_g1 | *F3'h* | 16.34 | 22.74 |
| RCL7_23161_c0_g1 | *F3'5'h* | 639.85 | 290.18 |
| RCL7_28129_c1_g1 | *Dfr1* | 610.03 | 77.41 |
| RCL7_20076_c4_g1 | *Dfr3* | 52.54 | 705.62 |
| RCL7_23829_c0_g1 | *Ans* | 579.14 | 35.97 |
| RCL7_20317_c4_g2 | *Uf3gt* | 189.3 | 107.65 |
| RCL7_23760_c0_g1 | *MYB6* | 664.67 | 354.33 |
| RCL7_25096_c3_g1 | *MYB11* | 117.44 | 36.33 |
| RCL7_27788_c0_g4 | *bHLH1* | 37.33 | 25.92 |
| RCL7_20277_c0_g2 | *bHLH2* | 1.85 | 1.52 |

| | | | |
|---|---|---|---|
| RCL7_22704_c0_g1 | *WDR1* | 19.48 | 11.32 |
| RCL7_22679_c0_g1 | *WDR2* | 10.62 | 7.38 |

Table S3. Voucher information for the *Clarkia* plants used in this study. RSABG, seeds obtained from Rancho Santa Ana Botanical Garden (Claremont, CA, USA). NFW, seeds from the collection of Norman F. Weeden. TRM, seeds provided by Talline R. Martins (University of Florida, Gainesville, FL, USA). LDG, seeds from the collection of Leslie D. Gottlieb.

| Specimen | Voucher | Location | Phenotype |
|---|---|---|---|
| *C. amoena huntiana* | RSABG 15875 | Mendocino County, CA | Central-spotted, pink-cupped |
| *C. amoena huntiana* | RSABG 15876 | Marin County, CA | Central-spotted, pink-cupped |
| *C. lassenensis* | NFW 84b | Shasta County, CA | Basal-spotted, white-banded |
| *C. lassenensis* | NFW 152 | Shasta County, CA | Basal-spotted, white-banded |
| *C. gracilis sonomensis* | TRM 14.6 (LDG 8513) | Sonoma County, CA | Central-spotted, pink-cupped |
| *C. gracilis sonomensis* | TRM 14.26 (LDG 8513) | Sonoma County, CA | Central-spotted, white-cupped |
| *C. gracilis albicaulis* | TRM 040 (Butte12) | Butte County, CA (Collected by B. Barringer) | Basal-spotted, white-banded |

**A**

```
CgsMYB1  ATGAATAAGGTAGGAGTTAGAAAGGGTGGTTGGACTGCAAATGAAGATGC  50
CgaMYB1  ATGAATAAGGTAGGACTTAGAAAGGGTGGTTGGACTGCAAATGAAGATGC  50
CahMYB1  ATGAATAAGGTAGGACTTAGAAAGGGTGGTTGGACTGCAAATGAAGATGC  50
 ClMYB1  -------------------------GTAGTTGGACAGCAAAAGAAGATGC  25

CgsMYB1  CCTACTCAAGCAATGCGTTCAAACTTATGGAGAAGGCAACTGGCATCTAG  100
CgaMYB1  CCTACTCAAGCAATGCATTCAAACTTATGGAGAAGGCAATTGGCATCTAG  100
CahMYB1  CCTACTCAAGCAATGCGTTCAAACTTATGGAGAAGGCAACTGGCATCTAG  100
 ClMYB1  CCTACTGAAGCAATGCGTTCAAACCTATGGAGAAGGTAACTGGCATCTAG  75

CgsMYB1  TCCCCGATAGAGCAGGTTTGAATAGGTGCAGAAAAAGTTGCCGATTAAGG  150
CgaMYB1  TCCCCGATCGAGCAGGTTTGAATAGGTGCAGAAAAAGTTGCCGATTAAGG  150
CahMYB1  TCCCCGATAGAGCAGGTTTGAATAGGTGCAGAAAAAGTTGCCGATTAAGG  150
 ClMYB1  TCCCCGATAGAGCAGGTTTGAATAGGTGCAGAAAAGGTTGCAGATTGAGG  125

CgsMYB1  TGGCTTAACTATCTGAAACCAGGCTTAAACCGAGAAGAGTTCCAAGAAGA  200
CgaMYB1  TGGCTTAACTATCTGAAACCAGGCTTAAACCGAGAAGAGTTCCAAGAAGA  200
CahMYB1  TGGCTTAACTATCTGAAGCCAGGCTTAAACCGAGAAGAGTTCCAAGAAGA  200
 ClMYB1  TGGCTTAACTATCTAAAGCCAGGCATAAATCGAGGAGAGTTCCAAGAAGA  175

CgsMYB1  TGAAATTGACTTGATTATTAGGCTTCACAAGCTTTTTGGCAATAAATGGT  250
CgaMYB1  TGAAATCGACTTGATTATTAGGCTTCACAAGCTTTTCGGCAAAAAATGGT  250
CahMYB1  TGAAATTGACTTGATTATTAGGCTTCACAAGCTTTTCGGCAAAAAATGGT  250
 ClMYB1  TGAAATTGACTTGATTATTAGGCTTCACAAGCTTTTTGGCAAAAAATGGT  225

CgsMYB1  CACTCATAGCAGGAAGACTTCCTGGAAGAACAAGCAATGATATAAAGAAT  300
CgaMYB1  CACTCATAGCAGGAAGACTTCCTGGAAGAACAAACAATGATATAAAAAAT  300
CahMYB1  CACTCATAGCAGGAAGACTTCCTGGAAGAACAAGCAATGATATAAAGAAT  300
 ClMYB1  CACTCATAGCAGGAAGACTTCCCGGAAGAACAAGCAACGATATAAAGAAT  275

CgsMYB1  TACTGGTACACCCATATTGCCAAGAAATTA---------CCTGCAGAACC  341
CgaMYB1  TACTGGTACACCCATATTGCCAAGAAATTA---------CCTGCAGAACC  341
CahMYB1  TACTGGTACACCCATATTGCCAAGAAATTA---------CATGCAGAACC  341
 ClMYB1  TACTGGAATAGTCATATTGTCAAGAAATTAGCCAAAGCTGCTGCACAACC  325

CgsMYB1  AGTTATATCGCAGGAAAATTCAGTCAAACACCACGCCATAATAAGGCCTA  391
CgaMYB1  AATTATATCGCAGGAAAATTCAGTCAAACACCACGCCATAATAAGGCCTA  391
CahMYB1  AATTATATCGCAGGAAAATTCAGTCAAACACCACGCCATAATAAGGCCTA  391
 ClMYB1  AATTATTTCGCAGGAAAATTCAGTCAAAGAACACGTCATAATAAAACCTA  375

CgsMYB1  TTGCAGGAAGACCGACCAAAGGGATGCAGTTTGGTATGAA----------  431
CgaMYB1  TTGCAGGAAGACCTACCAAAGGCATGCAGTTTGGTATGAA----------  431
CahMYB1  TTGCAGGAAGACCGACCAAAGGGATGCAGTTTGGTATGAA----------  431
 ClMYB1  TTGCAGGAAGACTTACCAAAGGGATGAAGTTTGGCATGAATATCGTCTTA  425

CgsMYB1  --TGTCACCTCTGATCAGCCTCCACCACCTCTGGAGAATTGGTCGGACTG  479
CgaMYB1  --TGTCACCTCTGATCAGCCTCCACCACCTCTGGAGAATTGGTCGGACTG  479
CahMYB1  --TGTCACCTCTGATCAGCCTCCACCACCTCTGGAGAATTGGTCGGACTG  479
 ClMYB1  GGTGCCACCTCTGATCAGCCTCCACCACCTTGGGAGAATTGGTGGGACAG  475

CgsMYB1  GTTGATGATGGACGATGATATTAATTATTATGATAATGGTGGTGGTTGTT  529
CgaMYB1  GTTGATGATGGACGATGATATTAATTATTATGACAATGGTGGTGGTTGTT  529
CahMYB1  GTTGATGATGGACGATGATATTAATTATTATGATAATGGTGGTGGTTGTT  529
 ClMYB1  GTTGATGATGGGTGATGATAATAATTATTATGATTTTGGTGGTGGTTGTT  525
```

Fig. S1. (to be continued on the next page)

```
CgsMYB1  CTGCCTCGGAAGGCCACTGCACCACCGCTGCCAATGGCTGCTACGACATG  579
CgaMYB1  CCGCCTCGGAAGGCCACTGCACCACCGCTGCCAATGGCTGCTACGACATG  579
CahMYB1  CTGCCTCGGAAGGCCACTGCACCACCGCTGCCAATGGCTGCTACGACATG  579
 ClMYB1  CCGCCTCAGGAGGCCACTGCACCACCGCTACCAATG---GCTACCACCTG  572

CgsMYB1  CAGATAGAATCTC---CGTGGGCGGCAATAGTAGGCGGATTCACCGAGGG  626
CgaMYB1  CAGATAGAATCTC---CGTGGGCGGCAATAGGAGGCGGATTCACCGAGGG  626
CahMYB1  CAGATAGAATCTC---CGTGGGCGGCAATAGTAGGCGGATTCACAGAGGG  626
 ClMYB1  CAGATAGGATCTCCGTCGTGTGCGGCAGAAGGA---------ACTGAGAG  613

CgsMYB1  TGGGAATAACCCGGATGAGCTTTATTTTGATGACATATTTTAA         669
CgaMYB1  TGGGAATAACCCGGATGAGCTTTATTTTGATGACATATTTTAA         669
CahMYB1  TGGGAATAACCCGGATGAGCTTTATTTTGATGACATATTTTAA         669
 ClMYB1  TGGGTATAACCC-------------------------------         625
```

Fig. S1. (to be continued on the next page)

**B**

```
CgsMYB6 ATGGGTGGTGTTCCTTGGACTGAAGAAGAGGATCTTCTGCTTAAGAAATG 50
CgaMYB6 ATGGGTGGTGTTCCTTGGACTGAAGAAGAGGATCTTCTGCTTAAGAAATG 50
CahMYB6 ATGGGTGGTGTTCCTTGGACTGAAGAAGAGGATCTTCTGCTTAAGAAATG 50
 ClMYB6 -------------------CTGAAGAAGAAGATCTTCTGCTTAAGAAATG 31

CgsMYB6 CGTCGAGCAGTTCGGCGAAGGGAAATGGCACCGGGTTCCGCTTTTAGCCG 100
CgaMYB6 CGTCGAGCAGTTCGGCGAAGGGAAATGGCACCGGGTTCCGCTTTTAGCCG 100
CahMYB6 CGTCGAGCAGTTCGGCGAAGGGAAATGGCACCGGGTTCCGCTTTTAGCCG 100
 ClMYB6 CGTCGAGCAGTTCGGTGAAGGGAAATGGCACCGGGTTCCGCTTTTAGCCG 81

CgsMYB6 GTCTGAACAGGTGCAGGAAGAGTTGCAGACTGAGGTGGCTGAATTATCTT 150
CgaMYB6 GTCTAAACAGGTGCAGGAAGAGTTGCAGACTGAGGTGGCTGAATTATCTT 150
CahMYB6 GTCTGAACAGGTGCAGGAAGAGTTGCAGACTGAGGTGGCTGAATTATCTT 150
 ClMYB6 GTCTAAACAGGTGCAGGAAGAGTTGCAGACTGAGGTGGCTGAACTACCTC 131

CgsMYB6 CGACCGAATATCAAGAGAGGGAGCTTTACTCAAGATGAAGTCGAGCTCAT 200
CgaMYB6 CGACCGAATATCAAGAGAGGGAGCTTCACTCAAGATGAAGTCGAGCTCAT 200
CahMYB6 AGACCGAATATCAAGAGAGGGAGCTTCACTCAAGATGAAGTCGAGCTCAT 200
 ClMYB6 CGACCGAATATCAAGAGAGGGAGCTTCACTCAAGATGAAGTCGAGCTCAT 181

CgsMYB6 CATCAAGCTCCATAAGCTTGTCGGGAATCGGTGGTCGATGATTGCCGGAA 250
CgaMYB6 CATCAAGCTCCATAAGCTTGTCGGGAATCGGTGGTCGATGATTGCCGGAA 250
CahMYB6 CATCAAGCTCCATAAGCTTGTCGGGAATCGGTGGTCAATGATTGCCGGAA 250
 ClMYB6 CATCAAGCTCCACAAGCTTGTCGGGAATCGGTGGTCGATGATTGCTGGAA 231

CgsMYB6 GACTCCCGGGAAGAACAGCTAACGATGTCAAGAACTTTTGGAACTGTCAT 300
CgaMYB6 GACTCCCTGGAAGAACAGCTAATGATGTCAAGAACTTCTGGAACTGTCAT 300
CahMYB6 GACTCCCGGGAAGAACAGCTAACGATGTCAAGAACTTTTGGAACTGTCAT 300
 ClMYB6 GACTCCCGGGAAGAACAGCGAACGATGTCAAGAACTTTTGGAACTGCCAT 281

CgsMYB6 CTAAGCAAAAAGCTGACTGCCGAACAAATGAGCATCGACCCTGAACAGAG 350
CgaMYB6 CTAAGCAAAAAGCTGACTGCCGAACAAATGAGCATCGACCCTGAACGAAG 350
CahMYB6 CTAAGCAAAAAGCTGACTGCCGAACAAATGAGCATCGACCCTGAACAGAG 350
 ClMYB6 CTGAGCAAAAAGCTGACTGCCGAACAAATGAGCATCGACCCTGAACGGAG 331

CgsMYB6 AATAGACAACCTAGTCCCGATAATAATGCCCCAACCGCGAAACCCTACAT 400
CgaMYB6 CATAGATAGCCTAGTCCCGGTCATAATGCCACAACCGCGAAACCCTGGAT 400
CahMYB6 AATAGACAACCTAGTCCCGATCATAATGCCCCAACCACGAAACCCTACAT 400
 ClMYB6 CATAGACAGTCTAGTCCCGGTCATAATGGCACAACCGCTAAAATCGGTAT 381

CgsMYB6 CCGTCTCAAGAAAACCGAACAAAAGAGACCAAGAAGTTGGGACTTCTATA 450
CgaMYB6 CCATCTCGAGAAAACCGAACAAAAGAGACCAAGAAGTTGGTACCTCTATA 450
CahMYB6 CCGTCTCGAGAAAACCGAACAAAAGAGACCAAGAAGTTGGGACTTCTATA 450
 ClMYB6 CCGTCTCGAGAAAACTGAACAAAAGAGAGCAAGAAGTTGGGACTTCTATA 431

CgsMYB6 GTGACACTTCCAAGTGTCGGTGAAGATGGAGCAATGAACGCAGTTCAAGT 500
CgaMYB6 GTGACACTTCCAAGTGTCGGTGAAGATGGAGCAATGAACGCAGTTCAAGT 500
CahMYB6 GTGACACTTCCAAGTGTCGGTGAAGATGGAGCAATGAACGCAGTTCAAGT 500
 ClMYB6 GTGACACTTCCAAGCGTCGGTGAAGATGGGGCAATGAATGCAATTCAAGT 481

CgsMYB6 TATCGATGATGGAAAGACGAACCCACCGAATCAAGAACACAGTGTCAGTC 550
CgaMYB6 TATCGATGATGGAAAGACAAACCCACCGAATCAAGAACACGGTGTCGGTC 550
CahMYB6 TATCGATGATGGAAAGACGAACCCACCAAATCAAGAACACGGTGTCAGTC 550
 ClMYB6 TATCGATCATGGAAAGATGAACCCTGCAACTCAAGAACACGGTGTCGGTC 531
```

Fig. S1. (to be continued on the next page)

```
CgsMYB6 TTGGAGATCTTCCCGGTGAATTCCAGTTCGACGAATGTAGATTAGACGGG 600
CgaMYB6 TTGGAGATCTTCCAGGTGAATTCCAGTTCGATGAATGTAGATTAGACGGG 600
CahMYB6 TTGGAGATCTTGCAGGTGAATTCCAGTTCGATGAATGTAGATTAGACGGG 600
 ClMYB6 TTGGAGATATTTCGGGAGAATTCCAGTTCGACGAATGTAGATTAGACGGG 581

CgsMYB6 ATTAGCAGCAGCAACAGCAGGAAATGGGACTGGGACGATCTGCTTATGGA 650
CgaMYB6 ATTAGCTGCAGCAACAGCAGGAAATGGGACTGGGACGATTTGCTCATGGA 650
CahMYB6 ATTAGCAGCAGCAACAGCAGGAAATGGGACTGGGACGATCTGCTCATGGA 650
 ClMYB6 ATAAGCAGCAGCAACAGCAGAAAATGGGATTGGGATGATCTACTCATGGA 631

CgsMYB6 TATGGATATCGATCTGTGGAATAACTCTCTGTAA                 684
CgaMYB6 TATGGATATCGATCTGTGGAATAACTCTCTGTAA                 684
CahMYB6 TATGGATATCGATCTGTGGAATAACTCTCTGTAA                 684
 ClMYB6 CATGGATAT-------------------------                 640
```

Fig. S1. (to be continued on the next page)

**C**

```
CgsMYB11 ATGAAGGGAGAATTAAGGAAGGGTGTTTGGAATGCAGAAGAAGATGCTCT 50
CgaMYB11 ATGAAGGGAGAATTAAGGAAGGGTGCTTGGAATGCAGAAGAAGATGCTCT 50
CahMYB11 ATGAAGGGAGAATTAAGGAAGGGTGCTTGGAATGCAGAAGAAGATGCTCT 50
 ClMYB11 ---------------------GGTGCATGGAATGCAGAAGAAGATGCTCT 29

CgsMYB11 CCTCAAGCAATGCATTCAAACTTATGGAGAAGGAAAATGGCATCTTGTTC 100
CgaMYB11 CCTCAAGCAATGCATTCAAACTTATGGAGAAGGAAAATGGCATCTTGTTC 100
CahMYB11 CCTCAAGCAATGCATTCAAACTTATGGAGAAGGAAAATGGCATCTTGTTC 100
 ClMYB11 CCTCAAGCAATGCATTCAAACTTATGGAGAAGGCAAATGGTATCTTGTTC 79

CgsMYB11 CTGCAAGAACAGGACTCAATAGGTGCAGAAAAAGTTGCAGGTTGAGGTGG 150
CgaMYB11 CTACAAGAACAGGACTCAATAGGTGCAGAAAAAGTTGCAGGTTGAGGTGG 150
CahMYB11 CTGCAAGAACAGGACTCAATAGGTGCAGAAAAAGTTGCAGGTTGAGGTGG 150
 ClMYB11 CCGCTAGAACAGGGCTGAATAGGTGCAGAAAAAGTTGCAGATTGAGGTGG 129

CgsMYB11 CTCAACTATCTGAAGCCGGGCATAAACCGTAGAGAGTTTCAAGAAGATGA 200
CgaMYB11 CTCAACTATCTGAAGCCGGGCATAAACCGTAGAGAGCTTCAAGAAGATGA 200
CahMYB11 CTCAACTATCTGAAGCCGGGCATAAACCGTAGAGAGCTTCAAGAAGATGA 200
 ClMYB11 CTCAACTATCTTAAACCGGGCATAAACCGTAAAGAGCTTCAAGAAGATGA 179

CgsMYB11 AGTTGACTTGATCATCAGGCTTCATAAGCTTCTTGGCAATAGATGGTCAC 250
CgaMYB11 AGTTGACTTGGTCATCAGGCTTCATAAGCTTCTTGGCAATAGATGGTCAC 250
CahMYB11 AGTTGACTTGATCATCAGGCTTCATAAGCTTCTTGGCAATAGATGGTCAC 250
 ClMYB11 AGTTGACTTGATCATCAGGCTTCATAAGCTTCTTGGAAATAGATGGTCAC 229

CgsMYB11 TTATTGCAGGAAGGCTACCGGGAAGAACATCCACCCAAGTAAAGAATTAC 300
CgaMYB11 TTATTGCGGGAAGGCTTCCGGGAAGAACATCCACCCAAGTAAAGAATTAC 300
CahMYB11 TTATTGCAGGAAGGCTTCCGGGAAGAACATCCACCCAAGTAAAGAATTAC 300
 ClMYB11 TTATTGCAGGAAGACTTCCCGGAAGAACATCCACCCACGTAAAGAATTAC 279

CgsMYB11 TGGAATGCCCATATAGCTAAAAAGTGGAGATCGTCATCCAAAGCTGCACC 350
CgaMYB11 TGGAATGCCCGTACAGCTAAAAAGTGGAGATCGTCATCCAAATCTGCAGC 350
CahMYB11 TGGAATGCCCATATAGCTAAAAAGTGGAGATCGTCATCCAAAGCTGCAGC 350
 ClMYB11 TGGAATGCCCATATAGCTAAAAAGTGGAGATCGTCATCAAAA------AGT 323

CgsMYB11 TGCAGAATCAAAATCATCATCTTATAAAGGAA---AACAGCAAGAATATT 397
CgaMYB11 TGCAGAATCAAAATCATCATC---TAAAGGAA---AACAACAACAATATT 394
CahMYB11 TGCAGAATCAAAATCATCATC---TAAAGGAA---AACAGCAAGAATATT 394
 ClMYB11 TGCAGAATCAAAATCATCATC---AAAAGGAAAGCAGCAGCAAAAATATC 370

CgsMYB11 CAGTAAACGTCATAAAGCCTATTGCTAGAAGAGCTCCCAAAATGATCGAT 447
CgaMYB11 CAGTAAACGTTATAAGGCCTATTGCTAGAAGAGCTCCCAAAATGATCGAT 444
CahMYB11 CAGTAAACGTCATAAAGCCTATTGCTAGAAGAGCTCCCAAAATGATCGAT 444
 ClMYB11 CAGTCAACGTCATAAGGCCTATTGCTAGAAGAGCTCCCAAAATGATCAAG 420

CgsMYB11 TTTGGTATGACCATGAATAACAATAATATATGCAGCATGTCCACCTCTCA 497
CgaMYB11 TTTGGTATGATCATGAATAACAATAATATATGCAGCAGGTCCACCTCTCA 494
CahMYB11 TTTGGTATGATCATGAATAACAATAATATATGCAGCATGTCCACCTCTCA 494
 ClMYB11 TTTGGT------ATGAATGATAATAATAAATGCAGCAGGTCCACCTCTCA 464

CgsMYB11 GCTGCCGC---------CTCATCTAGATTGTACCGGAATAATAAATACAG 538
CgaMYB11 GCTGCCGC---------CTCATCTAGATTGTACCGAAATAATAAATACAG 535
CahMYB11 GCTGCCGC---------CTCATCTAGATTGTACCGAAATAATAAATACAG 535
 ClMYB11 GCCGCCGCCGCCTCCATCTCATCTTGATTGTAACGAAATAATAAATGAAG 514
```

Fig. S1. (to be continued on the next page)

```
CgsMYB11  ACAAGGAGGTGATGAATTGGTTGCAGAGAATTATTGGACGATGATGATTTA  588
CgaMYB11  ACAAGGAGGTGATGAATTGGTTGGAGAGAATTATTGGACGATGATGATTTA  585
CahMYB11  ACAAGGAGGTGATGAATTGGTTGCAGAGAATTATTGGACGATGATGATTTA  585
 ClMYB11  ACAACGAGGTGATGAATTGGTTGGAGAGGTTATTGGACGATGATGATTTA  564

CgsMYB11  ATTGGTTTTGGAGGAGACGGTGGCTCTGCCGCCTCAGAAGGCCACTGTAC  638
CgaMYB11  ATTGGTGGTGGAGGAGACGGTGGCTCTGCCGCCTCAGAAGGCCACTGTAC  635
CahMYB11  ATTGGTGGTGGAGGAGACGGTGGCTCTGCCGCCTCAGAAGGCCACTGTAC  635
 ClMYB11  ATTGGTGGAGGAGGAGACAGTGGCTGTGCCGCCTCAGAAGGCCACTGTAT  614

CgsMYB11  CACCGTAGGTGGTGGGTATATGTCGACTCCATGA                  672
CgaMYB11  CACCGTAGGTGGTGGGTATATGTCGACTCCATGA                  669
CahMYB11  CACCGTAGGTGGTGGGTACATGTCGACTCCATGA                  669
 ClMYB11  CGTCGTAGAAGGTGGGTATATATCGACTGCATGA                  648
```

Fig. S1. (to be continued on the next page)

**D**

```
CgsMYB12  ATGAAGGAAGGTCTAAGGAAGGGTGCTTGGAGTGCAGAAGAAGATGCTCT  50
CahMYB12  --------------------GGGTGCTTGGAGTGCAGAAGAAGATGCTCT  30
 ClMYB12  ATGAACGGAGGTCTAAGGAAGGGTGCTTGGAGTGAAGAAGAAGATGCTCT  50

CgsMYB12  CCTCAAGCAATGTATTCAAATTTATGGAGAAGGCAAATGGCATCTTGTTC  100
CahMYB12  CCTCAAGCAATGTATTCAAATTTATGGAGAAGGCAAATGGCATATTGTTC  80
 ClMYB12  ACTCAGGCAATGCATTCAAACTTATGGAGAAGGCAAATGGCATCTTGTTC  100

CgsMYB12  CCGCCAGAGCAGGGCTAAATAGGTGTAGAAAAGGTTGCAGATTGAGGTGG  150
CahMYB12  CCGCCAGAGCAGGGCTAAATAGGTGTAGAAAAGGTTGCAGATTGAGGTGG  130
 ClMYB12  CCGCTAGATCAGGGCTGAATAGGTGCAGAAAAGGTTGCAGGTTGAGGTGG  150

CgsMYB12  CTCAACTATCTGAAGCCAGGCATAAACCTAAAAGAGCTTCAAGATGATGA  200
CahMYB12  CTCAACTATCTGAAGCCAGGCATAAACCTAAAAGAGCTTCAAGATGATGA  180
 ClMYB12  CTCAACTATTTGAAGCCCGGCATAAACCTAACAGAGCTTCAACATGATGA  200

CgsMYB12  AGTTGACTTGATCCTCAAACTTCACAAGCTTCTTGGCAACAAATGGTCAC  250
CahMYB12  AGTTGACTTGATCCTCAAACTTCACAAGCTTCTTGGCAACAAATGGTCAC  230
 ClMYB12  ACTTGACTTGATCTTCAAACTTCACAAGCTTCTTGGCAACAAATGGTCAC  250

CgsMYB12  TTATAGCAGGAAGACTTCCGGGAAGAACATGCAATTATATAAAGAATTAC  300
CahMYB12  TTATAGCAGGAAGACTTCCAGGAAGAACATGCAATTATATAAAGAATTAC  280
 ClMYB12  TTATTGCAGGAAGACTTCCCGGAAGAACATGCAACTACATAAAGAATTAC  300

CgsMYB12  TGGAACTCCAATATTGCTGCTAAAAAGTGGAAATCAAGAGAAAAGCAGCA  350
CahMYB12  TGGAACTCCAATATTGCTGCTAAAAAGTGGAAATCAAGAGAAAAGCAGCA  330
 ClMYB12  TGGAACTCCCATTTTACTGCTAAAAAGTGTAAATCAAGAAAAAAGCTGAA  350

CgsMYB12  AGATTCGGTCAAAGTTAAAGCCATAAGGCCTATTGTACGAAGAGCTCCCA  400
CahMYB12  AGATTCGGTCAAAGTCAAAGCCATAAGGCCTATTGTACGAAGAGCTCCCA  380
 ClMYB12  CGATTCGGTCAAAGT------CATAAGGCCTATTGTACGAAGAGCTCATA  394

CgsMYB12  AAATAATCAACTTTGGAATGAATAATAATAAT-------ATA--GGAACC  441
CahMYB12  AAATGATCAACTTTGGAATGAATAATAATAATAATAACAATAATGGAAGC  430
 ClMYB12  AAATGATCAAGTTCGGTAATAATAATAATAATAATAAC-ATA--GGGACC  441

CgsMYB12  ACCTCTCA---------------------------GCCTCATTGTAACGG  464
CahMYB12  ACCTCTCA---------------------------GCCTCATTGTAACGG  453
 ClMYB12  ACCTCTCGGCCGCCGTCACTGCCGCCAACACCTCCGCCTCATTGTAACGG  491

CgsMYB12  AATAACTAGGGACGACGAGGTGATGAATTGGTTGGATAGGTTACTAATGG  514
CahMYB12  AATAACTACAGACGACGAGGTGATGAATTGGTTGGATAGGTTACTAATGG  503
 ClMYB12  AATAAATACAGACAACGAGGTGGTGAATTGGTTGGATAGGTTACTAGTGG  541

CgsMYB12  ACGATGATGATGATGTTTATGCTTTTTGTAAAGGAGACGGGGGCTGTACC  564
CahMYB12  AC---GATGATGATGTTTATGCTTTTTGTAAAGGAGACGGTGGCTGTACC  550
 ClMYB12  AC------GATGATGTTTATGGTTTTGTTGGAGGAGACAGTGGTTGTTCC  585

CgsMYB12  GCCTCACAAGGCCACTGCTCCACCGCAGGTGGTGGCGGGTGTATCGACGG  614
CahMYB12  GCCTCACAAGGCCACTGCTCCACCGCAGGTGGT---GGGTGTATCGACAG  597
 ClMYB12  GCCTCACAAGGCCACTGTGCCATCGCAGGAGGT---GGGTATATCGACGA  632

CgsMYB12  CAGTGTCCTAGACGAGCTGTATATTGATCACGACATATTTCAGTTGTAG   663
CahMYB12  CAATGTCCTAGACGAGTTTTATATT------------------------   622
 ClMYB12  CAGTGTCCTAGACATGCTTTATATTGATCATCACATATTTCAGCTGTAG   681
```

Fig. S1. Alignment of *Clarkia R2R3-MYB* sequences (*A*) *MYB1*, (*B*) *MYB6*, (*C*) *MYB11* and (*D*) *MYB12* from *C. gracilis sonomensis*, *C. g. albicaulis*, *C. amoena* *huntiana* and *C. lassenensis*. If the nucleotides at each column are not identical, this column is highlighted in a black background. The sequences retrieved from GenBank are: *CgsMYB1* (KX592432), *CgaMYB1* (KX592431), *ClMYB1* (KX592428), *CgsMYB6* (MT425534), *CgsMYB11* (MT425536) and *CgsMYB12* (MT425538). Other sequences were generated in this study and were deposited in GenBank under the accession numbers MT796894-MT796902.

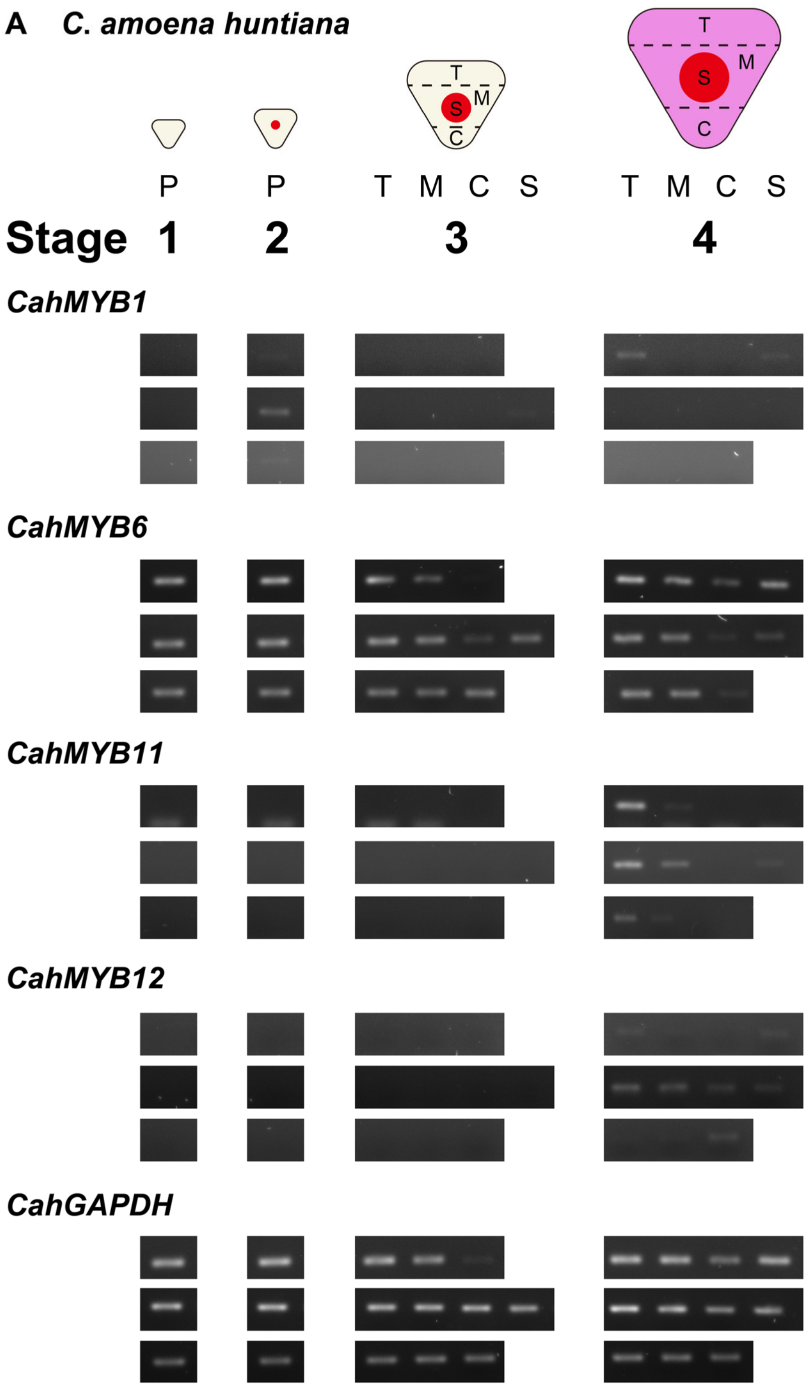


Fig. S2. (to be continued on the next page)

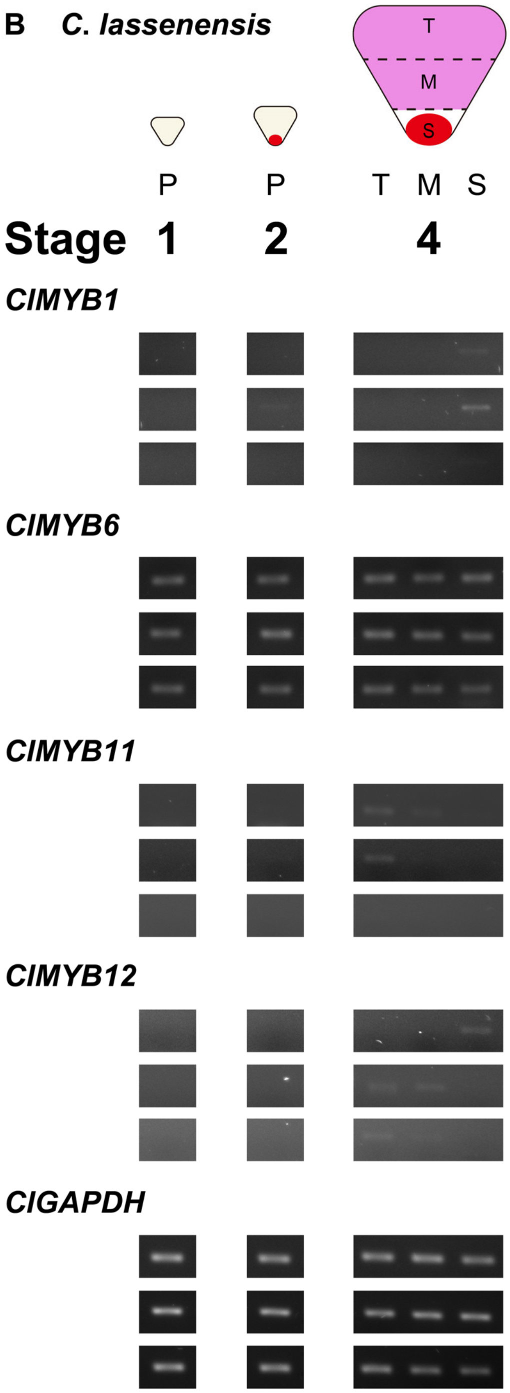


Fig. S2. (to be continued on the next page)

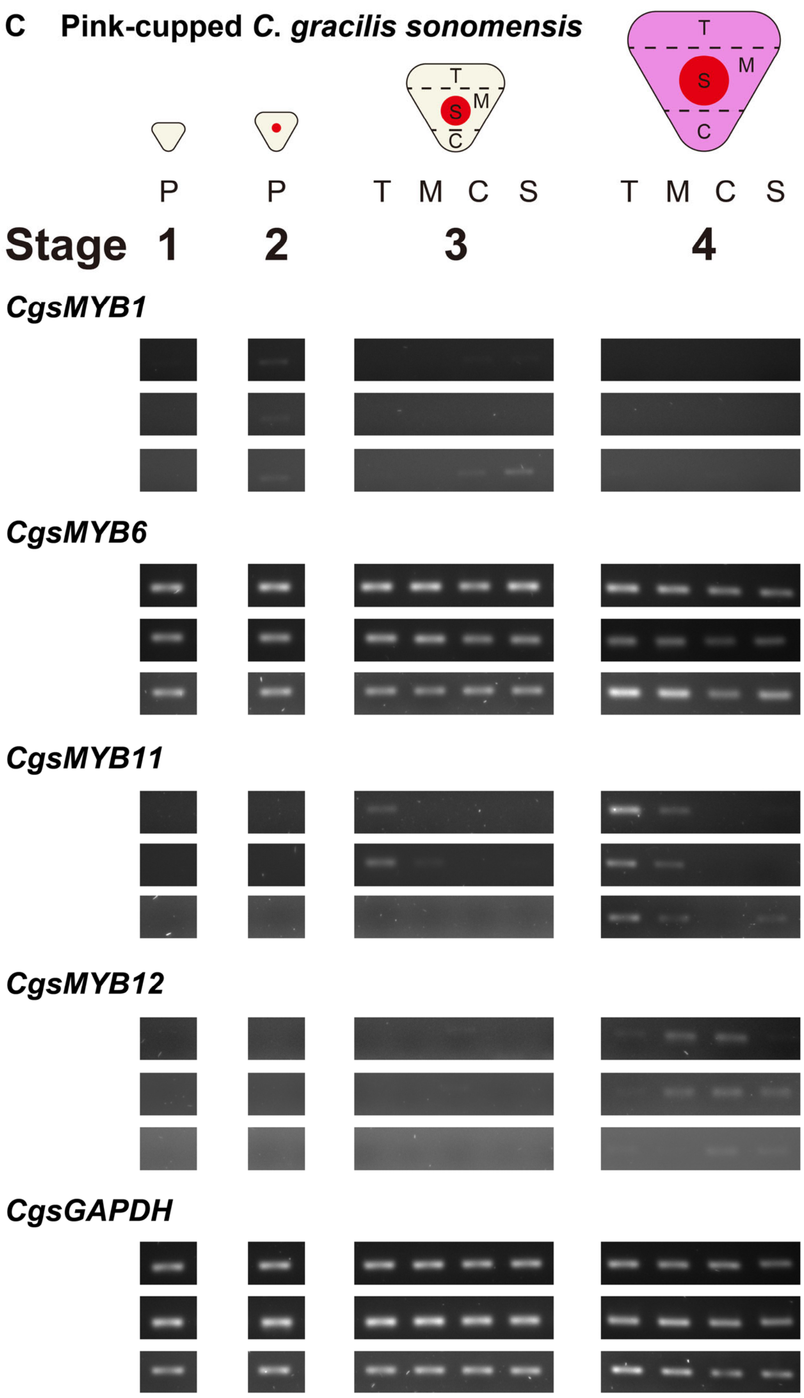


Fig. S2. (to be continued on the next page)

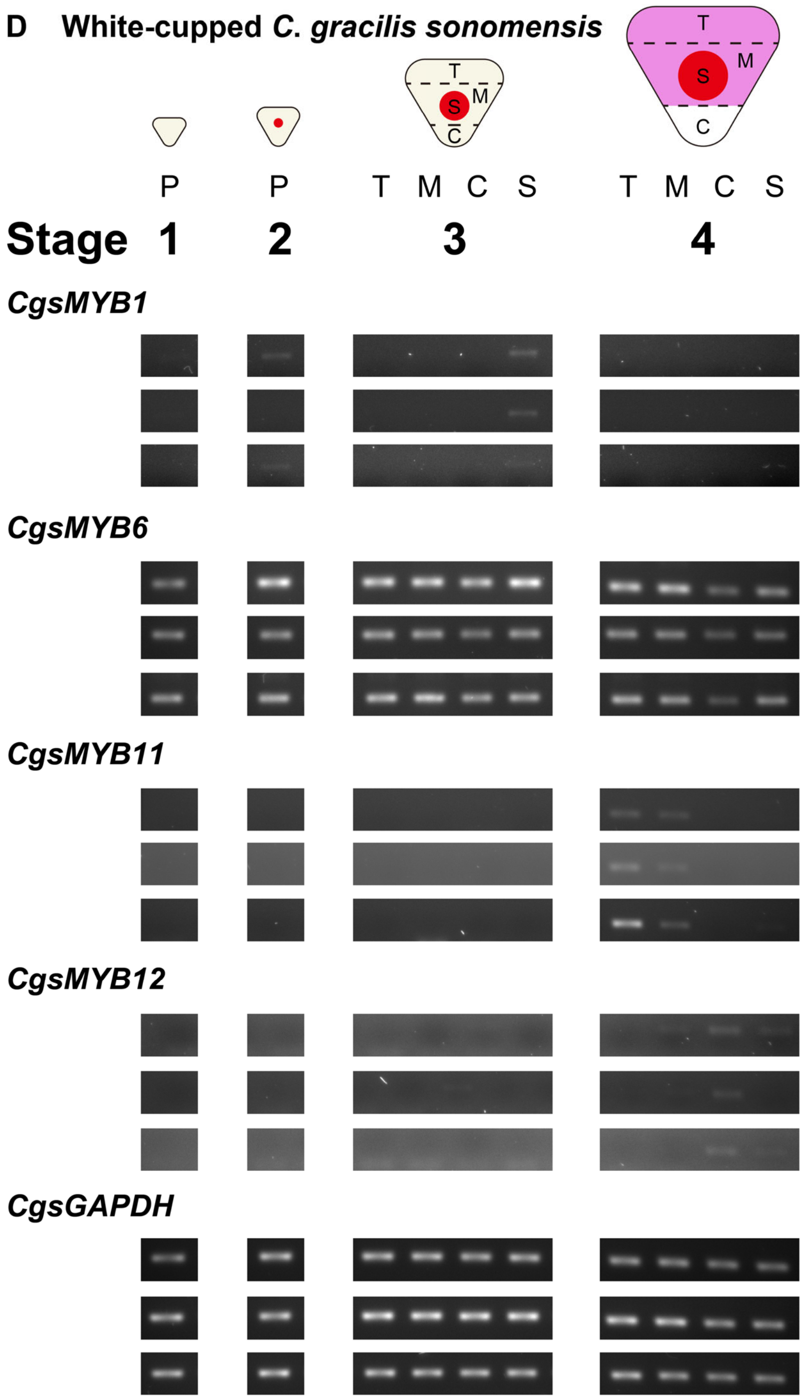


Fig. S2. (to be continued on the next page)

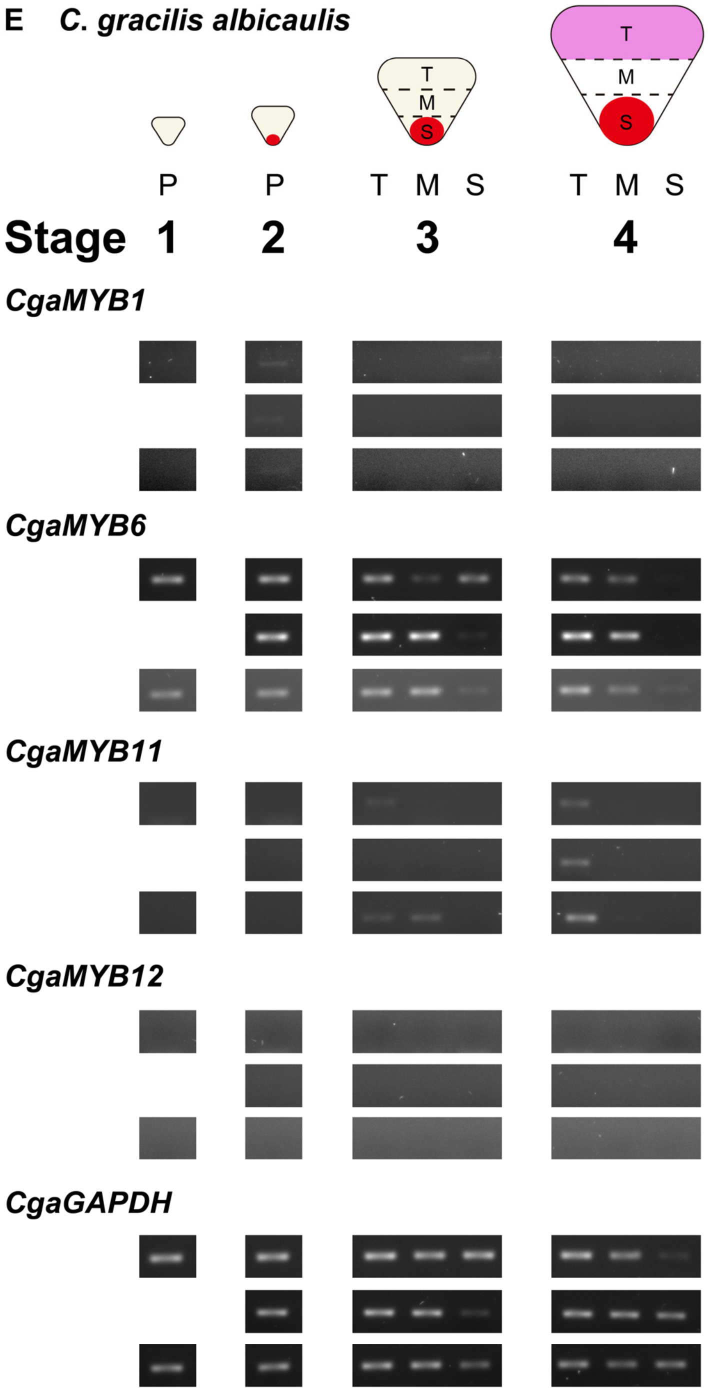


Fig. S2. Spatiotemporal expression of *MYB1*, *MYB6*, *MYB11* and *MYB12* in the petals of (*A*) *Clarkia amoena huntiana*, (*B*) *C. lassenensis*, (*C*) pink-cupped *C. gracilis sonomensis*, (*D*) white-cupped *C. g. sonomensis*, (*E*) *C. g. albicaulis*. Flower buds were collected from three plants from each (sub)species/phenotypes. For Stages 1 and 2, the whole petals (P) were used. For Stages 3 and 4, the petals were dissected into sections according to coloration: T (top), M (middle, excluding the spot), C (cup) and

S (spot). For *C. lassenensis*, Stages 2 and 3 were combined into Stage 2 due to a small petal size. A constitutively expressed gene *GAPDH* was included for cDNA quality control.

### Methods S1. Transcriptomics

To perform a comprehensive survey on which transcripts were expressed differentially between the pink background (the most distal portion) and the white band (the central portion) of the *Clarkia gracilis albicaulis* petals, we sequenced RNA extracted from these two petal regions. The petal regions were collected by dissecting flower buds (approximately 1 day before flowering) from four plants. Total RNA of pink background and white band was individually extracted using Spectrum Plant Total RNA Kit (Sigma-Aldrich, St. Louis, MO, USA). RNA samples of the two petal regions for library construction were prepared by pooling equal amounts of RNA from each plant. Prior to library construction, RNA quality was examined using Bioanalyzer Agilent RNA 6000 Nano Kit (Agilent Technologies, Santa Clara, CA, USA). We followed the protocol described in Supporting Information Methods S2 in Lin and Rausher (2021) to construct, barcode and sequence the libraries.

Bioinformatic analyses, including transcriptome assembling, identification of candidate genes for anthocyanin production and estimation of gene expression, were conducted as described in Lin and Rausher (2021).

Raw sequence reads generated in this study were deposited under NCBI Bioproject PRJNA721169 (Sequence Read Archive accessions: SRR14226355 and SRR14226356). The assembled transcriptomes were deposited at the NCBI Transcriptome Shotgun Assembly under the accessions GJDS00000000 and GJDT00000000. FPKM data are available at http://doi.org/10.5281/zenodo.4699974.
